# Regular aerobic exercise is positively associated with hippocampal structure and function in young and middle-aged adults

**DOI:** 10.1101/2020.08.14.250688

**Authors:** Joshua Hendrikse, Yann Chye, Sarah Thompson, Nigel C. Rogasch, Chao Suo, James Coxon, Murat Yücel

## Abstract

Regular exercise has numerous benefits for brain health, including the structure and function of the hippocampus. The hippocampus plays a critical role in memory function, and is altered in a number of psychiatric disorders associated with memory impairments (e.g. depression and schizophrenia), as well as healthy ageing. While many studies have focussed on how regular exercise may improve hippocampal integrity in older individuals, less is known about these effects in young to middle-aged adults. Therefore, we assessed the associations of regular exercise and cardiorespiratory fitness with hippocampal structure and function in these age groups. We recruited 40 healthy young to middle-aged adults, comprised of two groups (n = 20) who self-reported either high or low levels of exercise, according to World Health Organisation guidelines. We assessed cardiorespiratory fitness using a graded exercise test (VO2max) and hippocampal structure via manual tracing of T1-weighted magnetic resonance images. We also assessed hippocampal function using magnetic resonance spectroscopy to derive estimates of NAA concentration and hippocampal-dependent associative memory and pattern separation tasks. We observed evidence of increased N-acetyl-aspartate concentration and associative memory performance in individuals engaging in high levels of exercise. However, no differences in hippocampal volume or pattern separation capacity were observed between groups. Cardiorespiratory fitness was positively associated with hippocampal volume (left, right, and bilateral), N-acetyl-aspartate concentration, and pattern separation. However, no association was observed between cardiorespiratory fitness and associative memory. Therefore, we provide evidence that higher levels of exercise and cardiorespiratory fitness are associated with improved hippocampal structure and function. Exercise may provide a low-risk, effective method of improving hippocampal integrity in an early-to-mid-life stage.

## Introduction

A physically active lifestyle is associated with a myriad of systemic health benefits, including a reduced risk of type 2 diabetes, cardiovascular disease, and certain forms of cancer (Bassuk & Manson, 2005; McTiernan, Schwartz, Potter, & Bowen, 1999). It has become increasingly clear that the benefits of physical activity (PA) are not limited to the periphery, but also extend to the brain (Voss, Vivar, Kramer, & van Praag, 2013). PA refers to forms of bodily movement that result in energy expenditure, and epidemiological studies have associated higher engagement in PA with greater preservation of cognitive capacity (Yaffe, Barnes, Nevitt, Lui, & Covinsky, 2001), and reduced incidence of dementia in later life (Laurin, Verreault, Lindsay, MacPherson, & Rockwood, 2001; Rovio et al., 2005). A number of meta-analyses have also reported improved cognitive performance following exercise interventions (Angevaren, Aufdemkampe, Verhaar, Aleman, & Vanhees, 2008; Smith et al., 2010), particularly within the memory domain (Smith et al., 2010).

Exercise is a subcategory of PA and refers to any deliberate, repetitive movement intended to maintain or improve physical fitness. Aerobic forms of exercise increase respiration and heart rate to increase delivery of energy substrates to working skeletal muscle. Engaging in exercise (especially aerobic forms such as running) represents a practical method of improving cognitive performance and mitigating age-related cognitive decline, and may also offer an effective treatment alternative across a number of psychiatric and neurological disorders (Hillman, Erickson, & Kramer, 2008). However, to maximise clinical potential, a comprehensive understanding of the underlying neural mechanisms targeted by exercise is required.

The hippocampus plays a critical role in memory (Eichenbaum, Schoenbaum, Young, & Bunsey, 1996) and affective processing (Fanselow & Dong, 2010), and a number of studies have associated exercise with changes to hippocampal structure and function (Kandola, Hendrikse, Lucassen, & Yücel, 2016). For example, increased hippocampal volume has been observed following exercise interventions in older adults (Colcombe et al., 2006; Erickson et al., 2011; Niemann, Godde, & Voelcker-Rehage, 2014), and a positive association between self-reported weekly exercise and right hippocampal grey matter volume has been demonstrated in young to middle-aged adults (Killgore, Olson, & Weber, 2013). Furthermore, higher cardiorespiratory fitness (CRF) level, reflecting physiological adaptation in response to regular PA/exercise, is associated with larger hippocampal volume in preadolescents (Chaddock et al., 2010) and older adults (Colcombe et al., 2003). At a micro-scale level, exercise also has demonstrable benefit on the biochemical composition of the hippocampus. N-acetyl-aspartate (NAA) is primarily localised to the cell body of neurons and has a critical role in lipid synthesis and cellular metabolism, and is therefore regarded as a marker of neuronal integrity/vitality (Moffett, Ross, Arun, Madhavarao, & Namboodiri, 2007). Elevations in NAA concentration as measured by magnetic resonance spectroscopy (MRS) have been observed following aerobic exercise interventions in healthy adults (den Ouden et al., 2017) and individuals with schizophrenia (Pajonk et al., 2010). Overall, these studies suggest that exercise can influence hippocampal structure. However, the functional relevance of exercise-induced structural plasticity remains incompletely understood.

Exercise has been linked to enhancements of hippocampus-dependent forms of memory, such as declarative memory (Kandola et al., 2016). Declarative memory encompasses associative and episodic memory processes (referring to the recollection of relationships between cues and autobiographical events respectively), which are also inherently related to spatial memory (referring to spatial navigation/orientation in one’s environment) (Squire & Zola, 1996). Similarly, the hippocampus also supports the ability to select and distinguish similar spatial and temporal memory traces, a process referred to as ‘pattern separation’ (Kirwan & Stark, 2007; Stark, Yassa, Lacy, & Stark, 2013). Using cognitive tasks designed to assess these forms of memory, studies have shown positive associations between CRF and episodic/associative memory (Baym et al., 2014; Flöel et al., 2010) and pattern separation capacity following exercise interventions (Déry et al., 2013). However, while the evidence seems to support a role for exercise in the promotion of hippocampal function, a number of studies have failed to demonstrate beneficial effects (for review see, Kandola, Hendrikse, Lucassen, & Yücel, 2016). Further, while some studies have examined the effects of exercise on the hippocampus of young to middle-aged adult samples (e.g. Déry et al., 2013; Killgore et al., 2013; Pajonk, Wobrock, Gruber, & et al., 2010; Thomas et al., 2016b; Wagner et al., 2015), the majority of studies have been conducted on older adults, i.e. > 65 years of age (Prakash, Voss, Erickson, & Kramer, 2015; Stillman, Esteban-Cornejo, Brown, Bender, & Erickson, 2020). At present, there is insufficient evidence that exercise and cardiorespiratory fitness benefits hippocampal structure and function in young to mid-adulthood. For instance, it remains unclear whether engaging in regular exercise (e.g. meeting current American College of Sports Medicine (ACSM) / World Health Organisation (WHO) guidelines of at least 150 minutes moderate intensity aerobic activity each week) is associated with improved hippocampal structure and function in these age groups. Thus, an improved understanding of the relationship between exercise and hippocampal outcomes at this stage of life is required.

The aim of this study was to investigate associations of exercise and CRF with hippocampal structure and function in young to middle-aged adults. We recruited a sample of 40 healthy adults, comprised of individuals who self-reported as engaging in a high level of exercise (EXE) and met the WHO/ACSM guidelines (high EXE, n = 20) and low levels of exercise below the guidelines (low EXE, n = 20). A multi-modal assessment of hippocampal structure and function was conducted using structural magnetic resonance imaging (MRI), MRS, and hippocampal-dependent behavioural tasks. Specifically, we employed measures of associative memory (Wang et al., 2014) and pattern separation (Stark, Yassa, Lacy, & Stark, 2013) as it is unclear whether these aspects of hippocampal function are benefitted by regular exercise in these age groups. CRF was also quantified by having participants complete an incremental exercise test to measure of maximal oxygen consumption i.e. VO_2_max. We expected significant differences in hippocampal structure and function between high EXE and low EXE groups. Specifically, we expected higher hippocampal volume and NAA concentration in the high EXE group, relative to the low EXE group. Similarly, we expected that the high EXE group would demonstrate higher performance on associative memory and pattern separation tasks. We also hypothesised positive associations between CRF and measures of hippocampal structure and function (i.e. volume, NAA concentration, and pattern separation and associative memory performance).

## Methods

### Ethics approval

This study was approved by the Monash University Human Research Ethics Committee and all participants provided informed consent. The study conformed to the standards set by the Declaration of Helsinki, and participants were remunerated $100 AUD for their participation as part of a larger study.

### Participants

Participants were 40 right-handed healthy adults (52.5% female), aged 25.48 ± 9.35 years (mean ± SD; range 18 – 55), reporting no contraindications to MRI and no history of psychological or neurological disorders. To investigate associations between exercise and hippocampal structure and function, participants were recruited and selected on the basis of self-reported weekly exercise into a high EXE or low EXE group. The high EXE group included individuals engaging in a minimum of 150 minutes of deliberate exercise each week (mean weekly exercise 428.5 minutes ± 215.6; mean ± SD, range = 150 – 900 minutes), consisting primarily of sports/physical activities known to increase CRF level (e.g. running, swimming, and cycling). The low EXE group included individuals engaging in no more than 60 minutes of deliberate exercise each week (mean weekly EXE 13.8 minutes ± 25.0; mean ± SD, range = 0 – 60 minutes, Mann-Whitney test, p=3×10^−8^, r = 1.00). All participants were required to complete the long format of the International Physical Activity Questionnaire (IPAQ) (Craig et al., 2003) to verify self-reported exercise level prior to participation.

### Experimental design

A between-subjects, cross-sectional design was used to investigate associations between exercise and hippocampal structure and function. Participants completed an MRI scan (approximately 45 minutes in duration), followed by two assessments of memory function. Comparisons of hippocampal structure (left and right hippocampal volume and NAA concentration) and hippocampal-dependent memory tasks (face-word recall and mnemonic similarity task) were made between high and low EXE groups. In a separate session, participants completed a VO_2_max test to assess cardiorespiratory fitness level.

### Cardiorespiratory fitness assessment (VO_2_max)

To determine each subject’s CRF level, maximal oxygen consumption (VO_2_max) was assessed by a graded intensity maximal exercise test conducted on a motorised treadmill. VO_2_max was conducted to provide an objective measure of individual fitness to correlate with measures of hippocampal structure and function. The starting pace was set at 6km/h with a 1% gradient, and workload was increased every three minutes until volitional exhaustion was reached. This was achieved via increasing speed by 2 km/h until 16 km/h, followed by incremental gradient increase of 2.5%. Heart rate, expired air volume, and concentration of oxygen and carbon dioxide were continuously recorded throughout the test duration. For determination of VO_2_max on the basis of VO2peak, at least two of the following indicators needed to be observed: (1) a plateau in VO_2_ml/kg/min values irrespective of increased workload, (2) maximal respiratory exchange ratio of ≥ 1.1, (3) a heart rate within 10 beats of age-predicted heart rate max (Tanaka, Monahan, & Seals, 2001), or (4) a self-reported rating of perceived exertion (BORG scale; Borg, 1982) rating of perceived exertion > 17 out of 20. Three participants from the low EXE group did not meet these criteria and recorded a submaximal test outcome. For these participants, individual regression equations were used (Mean R^2^ = .77) to derive a predicted VO_2_max value on the basis of age-predicted max heart rate (Tanaka et al., 2001). VO_2_max is reported in millilitres of oxygen per minute per kilogram of body mass (ml/kg/min) to account for individual differences in body weight.

### Magnetic resonance imaging (MRI)

MRI data were collected from a Siemens 3T Skyra scanner with 32-channel head coil. T1-weighted structural images (Magnetization Prepared Rapid Gradient Echo, TR = 2.3 s, TE = 2.07 ms, voxel size 1mm^3^, flip angle 9°, 192 slices) and GABA-edited Mescher-Garwood Point Resolved Spectroscopy (MEGA-PRESS) data were acquired. MEGA-PRESS data were acquired from a 2 × 3 × 1.5cm voxel localised to the left hippocampus (96 ON-OFF averages, TE = 68 ms, TR = 1.5 s, editing pulses at 1.9 ppm and 7.5 ppm with bandwidth = 45 Hz). NAA concentration was estimated from the ‘OFF’ average spectra (see figure 1). Following each acquisition, 8 ON-OFF averages with unsuppressed water were also obtained using the same parameters.

**Figure 1.**
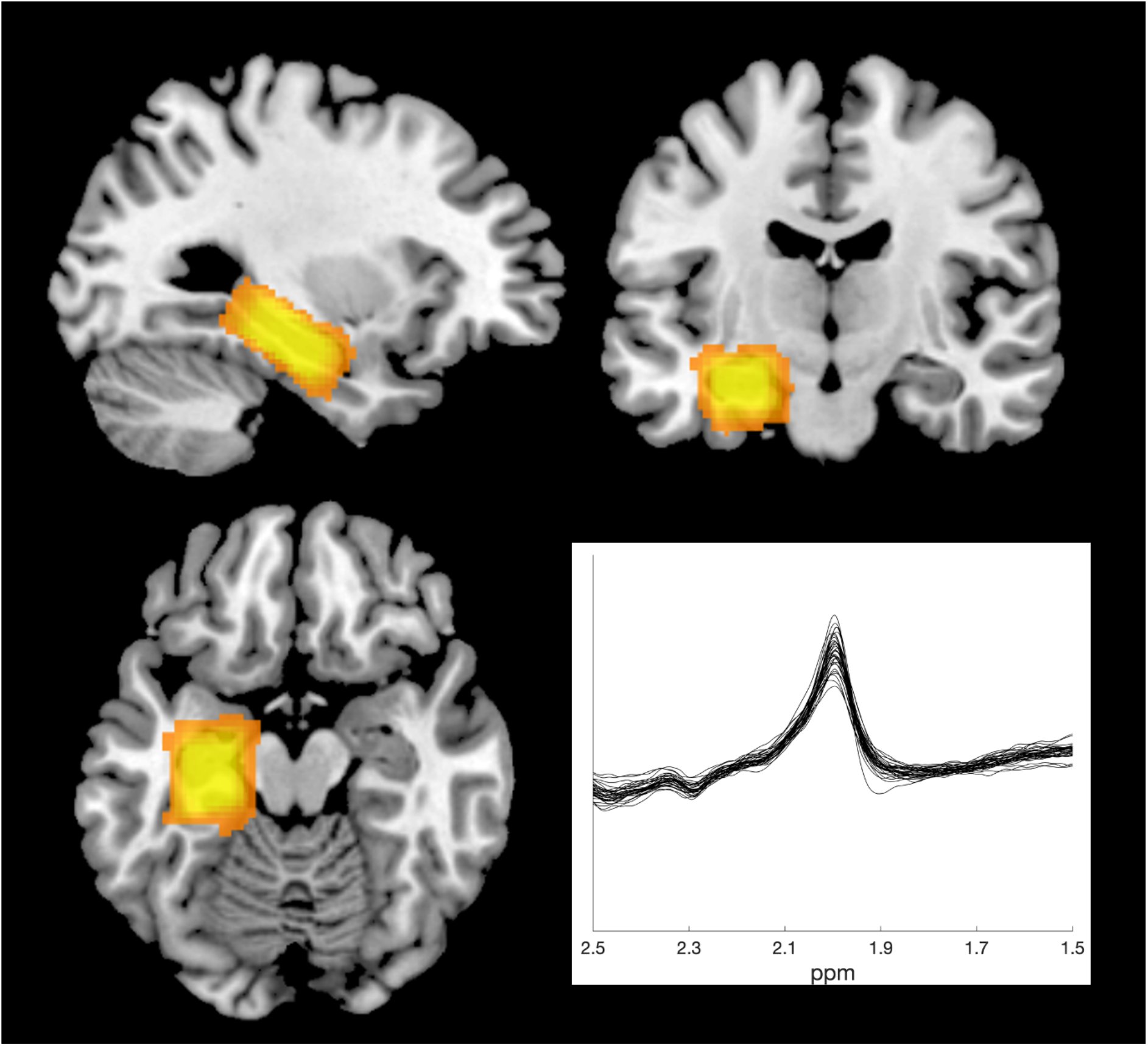
Positioning of MR spectroscopy voxel over left hippocampus and NAA spectra. MEGA-PRESS data were acquired from 2 × 3 × 1.5 cm voxels localised to left hippocampus (top row and bottom left). The overlap in voxel position across high and low EXE participants is shown in orange/yellow, with greater spatial overlap represented by lighter colouration. NAA concentration estimates were derived from the unedited ‘OFF’ spectra (bottom right; spectra shown for all participants with NAA peak occurring at 2.02 ppm) and reported relative to the unsuppressed water signal.

### Memory assessment

#### Associative memory – Face-cued word recall

To assess the effects of exercise on associative memory function, we employed a face-cued word recall task (Wang et al., 2014). Participants studied a set of 20 human face photographs derived from a database of amateur model headshots (Althoff & Cohen, 1999), presented in greyscale on printed cards. Each card was placed on a table in front of the participant for three seconds, and a unique common English word was read aloud by the experimenter when each card was shown. The words were nouns between 3-8 letters in length, with Kucera-Francis written frequencies between 200 to 2000, and concreteness ratings of 300 to 700 (MRC Psycholinguistic Database; www.psych.rl.ac.uk). Participants were instructed to memorise the association between the face and word. Following presentation of the 20 face-word pairs, there was a filled delay of approximately one-minute duration. Following this, the same set of cards was represented to participants individually in a different randomised order, and participants were instructed to try and recall the word that accompanied each face. Word recall was scored as correct or incorrect, forgiving errors relating to pronunciation, including plurality. Five alternative versions of the task were used across participant groups (matched for word concreteness and frequency) and randomised across participants using a Latin square. To familiarise participants with the task and minimise practice effects, a sixth version of the task was employed as a practice set, whereby a smaller sub-set of 10 face-word pairs was administered prior to the main assessment.

#### Pattern separation - mnemonic similarity task

To investigate differences in behavioural pattern separation between EXE groups, we used the mnemomic separation task developed by Stark, Yassa, Lacy, & Stark (2013). Performance on the task is correlated with increased BOLD activity in hippocampal sub-regions (Kirwan & Stark, 2007) and improvements have been reported following a six week aerobic exercise intervention (Déry et al., 2013). In the task, participants were required to complete an initial encoding phase in which they were presented with a series of 96 colour images of everyday objects in the centre of a screen on a white background. Participants were required to categorise each image as an ‘indoor’ or ‘outdoor’ item via button press, using first and third fingers of their right hand (V key for indoor items, N key for outdoor items). Each image was presented for 2.5 s with a 0.5 s inter-trial interval. After a short ~30 s delay, participants were given instructions about a surprise recognition test during which a second series of 96 colour images were presented (2.5 s each, 0.5 inter-trial interval). Participants were required to identify whether the image was old, similar, or new via button press using first three fingers of their right hand (V key for old, B key for similar, N key for new). One third of images were identical repetitions of previously seen images (i.e. old/targets), one third were new images that had not been previously presented (i.e. new/foils), and one third were similar but not identical to those previously presented (i.e. similar/lures). Responses to each trial during the recognition phase were coded as correct or incorrect.

### Data analysis

#### Exercise and cardiorespiratory fitness

We assessed differences in self-reported weekly exercise and CRF level (VO_2_max) between EXE groups. Two participants from the high EXE group were lost to follow-up and did not complete the graded exercise test. Thus, analyses of CRF level were conducted with N = 38 (high EXE, n = 18; low EXE, n = 20). We also assessed the association between VO_2_max and current self-reported weekly exercise using a Pearson’s correlation.

#### Analysis of hippocampal volume

Left and right hippocampi were manually traced by an expert tracer (Y. C.) (Chye et al., 2019) using each subject’s T1-weighted image and Analyze 12.0 software (AnalyzeDirect, Overland Park, KS) (see figure 2). Hippocampal tracing was conducted according to a validated protocol (Velakoulis et al., 1999), whereby the tracer was blinded to EXE group categorisation and left/right structural image orientation. Posterior hippocampal boundaries were defined as the slice with the greatest length of continuous fornix; medial boundaries as the open end of the hippocampal fissure (posterior) and the uncal fissure (anterior); lateral boundaries as the temporal horn of the lateral ventricle; inferior boundaries as the parahippocampal white matter; and superior boundaries as the fimbria and alveus (posterior) as well as the amygdala (anterior). To assess the intra-rater reliability of manual traced images, each subject’s hippocampi were retraced from a T1-weighted structural image acquired one week after the experimental assessment. There were no significant differences between hippocampal volume between time points (see supplementary figure S2). The intra-rater correlation coefficient between experimental and follow-up assessments was .89 for left hippocampus and .86 for right hippocampus, demonstrating consistent manual tracing of hippocampi (see supplementary materials for scatterplots). For each subject, total brain volume estimates were calculated using the ‘fsl_anat’ function (Jenkinson, Beckmann, Behrens, Woolrich, & Smith, 2012). Total brain volume was used to control for differences in brain size across participants and normalise hippocampal volume estimates.

**Figure 2.**
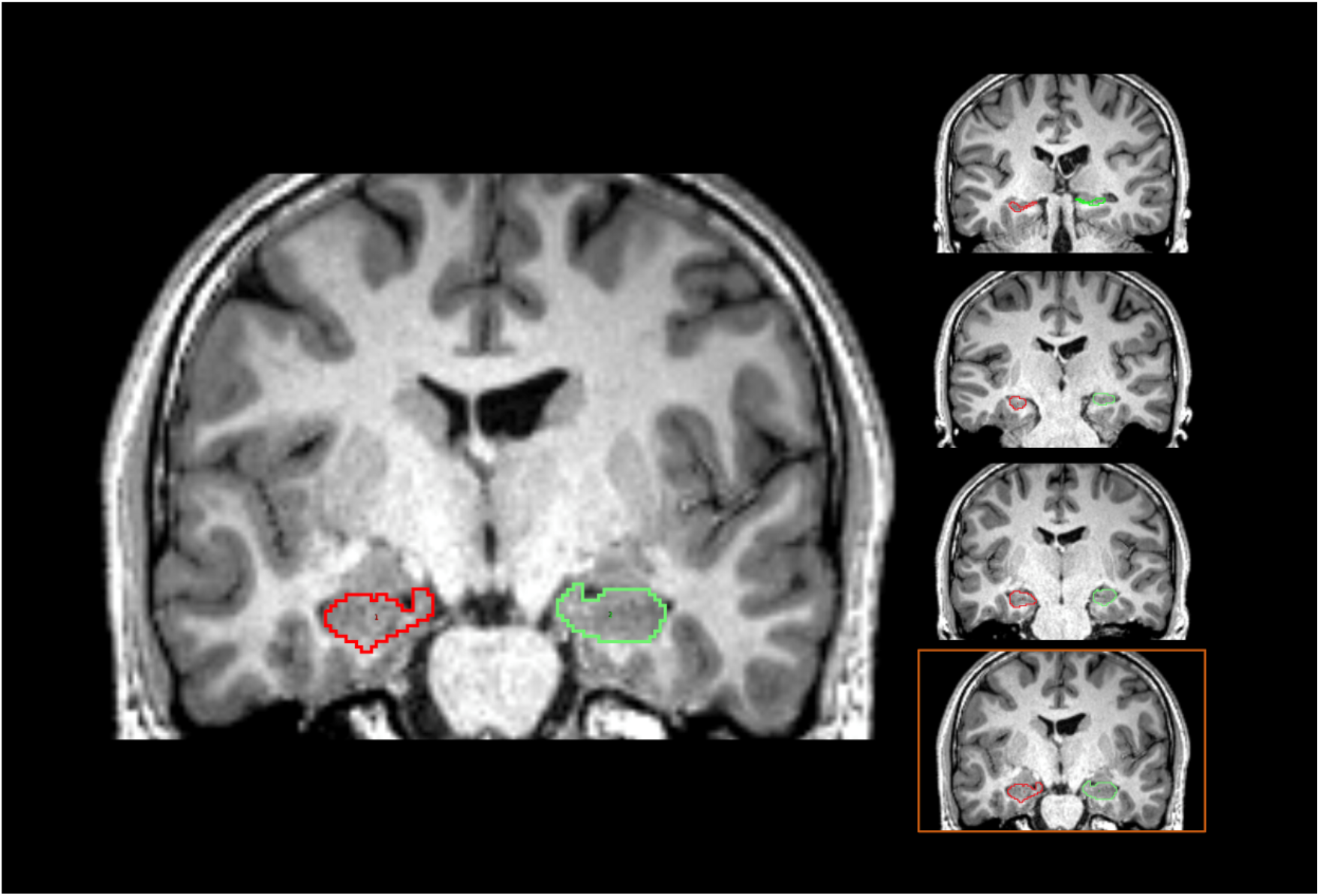
Manual tracing of the left and right hippocampus. Left and right hippocampi were manually traced by an expert tracer using each subject’s T1-weighted image. Tracing was completed for left (red outline) and right (green outline) hippocampi according to a validated protocol. Figure represents the manual tracing process, with hippocampi traced across each slice of the T1-weighted image in the coronal plane.

#### NAA quantification

MEGA-PRESS data were acquired as part of a wider study assessing cortical GABA concentration. Relative to cortical regions, the hippocampus is associated with a lower signal-to-noise ratio, reducing the reliability of MRS GABA estimates. Therefore, in this study, we focussed on NAA concentration via the unedited PRESS spectra. NAA is present in much higher concentrations than GABA and produces the largest peak in (unedited) MRS scans of the healthy human brain (Moffett, Arun, Ariyannur, & Namboodiri, 2013), and is therefore one of the most reliable MRS signals (Moffett et al., 2007). MRS data was not collected for two subjects due to scheduling issues, thus analyses were conducted with N = 38. Gannet (version 3) (Edden et al., 2014) was used to estimate NAA concentration from the left hippocampus. The GannetLoad module was employed for phased-array channel combination of raw Siemens ‘twix’ files, exponential line broadening (3 Hz), Fourier transformation to yield time-resolved frequency-domain spectra, frequency and phase correction, outlier rejection, and time averaging. GannetFit module was employed to estimate the area of the NAA peak from the unedited ‘OFF’ acquisition using a single gaussian model and a nonlinear least-squares fitting approach. As an internal reference, NAA concentrations are expressed in institutional units relative to the unsuppressed water signal, which was estimated with a Gaussian-Lorentzian model.

Partial volume segmentation of each subject’s T1-weighted anatomical image within each voxel was conducted using FSL’s FAST(Jenkinson et al., 2012), and NAA:H2O ratios were partial volume corrected by removing the contribution of cerebrospinal fluid fractions (NAA:H_2_O / (1 - CSF%)). To assess the quality of NAA estimates, we compared the full width at half maximum (FWHM) values of NAA and water peaks, water concentration, and NAA fit error (E_NAA_) between EXE groups. E_NAA_ is a measure reflecting the SD of model fit residual for the NAA peak, expressed as a percentage of peak amplitude. Spectra with an E_NAA_ value of > 20% or exceeding a z-score of 3.29 were labelled as outliers and removed from the analysis.

#### Face-cued word recall task

Performance on the face-cued word recall task was scored as the percentage of correctly recalled words out of a total of 20. Individual z scores > 2.58 were labelled as outliers and removed from subsequent analysis.

#### Mnemonic similarity task

Data from one subject (high EXE) was lost due to computer malfunction during data acquisition, thus analyses are reported with N = 39. Performance on the mnemonic similarity task was calculated by a separation bias metric, which indexed subject’s ability to discriminate ‘similar’ items from ‘new’ items, i.e. probability(respond ‘similar’ on ‘similar’ image) – probability(respond ‘similar given ‘new’ image), and also the overall percentage of correct responses.

#### General statistical approach

Statistical analysis of these data was conducted using JASP (v 0.10.2). An alpha level of .05 was adopted for all inferential statistics conducted under the null hypothesis significance testing framework. For all frequentist statistics, p-values should be interpreted in light of this .05 test-wise alpha level. This approach was complemented with measures of effect size, and Bayesian analysis to assess the relative evidence in favour of the alternative hypothesis (i.e. BF_10_). BF_10_ values between 1-3 were considered anecdotal evidence against the null, > 3 substantial evidence against the null, and < 0.33 substantial evidence for the null (Wagenmakers, Wetzels, Borsboom, & van der Maas, 2011).

#### Statistical comparisons between high and low EXE groups

To assess differences in CRF, self-reported exercise level, NAA: H_2_O ratios, and hippocampal function (face-cued word recall, pattern separation), independent-samples t-tests were conducted between high and low EXE groups (or non-parametric equivalent in the case of violations of normality). Left, right, and bilateral hippocampal total volume (mm^3^) were analysed in separate ANCOVA models, with a dependent factor of Hippocampal Volume, fixed factor of EXE Group (high EXE, low EXE), and Total Brain Volume (mm^3^) as a covariate. Bayesian forms of these tests were conducted to quantify the relative evidence for the alternative vs null hypothesis (priors set in support of the alternative over null hypothesis (i.e. BF_10_), and Cauchy parameter set to a conservative default value of 0.707 for independent-samples t-tests (Ly, Verhagen, & Wagenmakers, 2016; Rouder, Speckman, Sun, Morey, & Iverson, 2009).

#### Relationship between cardiorespiratory fitness and hippocampal structure and function

We assessed the relationship between CRF level and hippocampal structure (volume, NAA concentration) and function (i.e. associative memory, pattern separation). Upon inspection of the data, a relatively continuous distribution of VO_2_max scores was found across EXE groups (see figure 3). CRF/ VO_2_max provides an indirect measure of habitual exercise levels (Carrick-Ranson et al., 2014) and is also influenced by a number of factors such as age, biological sex, and genetic/physiological predispositions. Thus, the continuous distribution of VO_2_max scores across EXE groups is not particularly surprising, despite the deliberate selection of high and low EXE individuals. Instead, this highlights that CRF and self-reported weekly exercise are different exercise parameters and may uniquely explain hippocampal outcomes. Given the continuous distribution of CRF levels, multiple linear regression was used to examine the associations between CRF (VO_2_max) and hippocampal measures. To assess the relationship between VO_2_max and hippocampal volume, separate multiple linear regression models were conducted on left, right, and bilateral volume estimates, with age, sex, and total brain volume as additional covariates. Multiple linear regression was also used to examine the relationship between VO_2_max and NAA, with age and sex as additional covariates (see supplementary materials for additional control analysis, accounting for potential influence of grey and white matter tissue components). Similarly, multiple linear regression was used to examine the extent to which CRF was associated with hippocampal function (face-cued word recall and mnemonic discrimination performance), with age as an additional covariate. Biological sex was included as a covariate for analysis of hippocampal volume and NAA concentration based on evidence of sex-based differences in hippocampal morphology (Persson et al., 2014; Ruigrok et al., 2014) (but see Tan, Ma, Vira, Marwha, & Eliot, 2016). However, due to the paucity of evidence of sex-based differences in associative memory and pattern separation in humans, biological sex was not included as a covariate for these analyses (see supplementary materials for additional analyses with biological sex as a covariate in the regression model, and for comparisons of hippocampal function between male and female subjects). For each regression model, we calculated Cook’s distances (Cook, 1977) to assess the leverage/influence of individual values on the regression line of best fit. Individual values exceeding a Cook’s distance of three times the mean were labelled as outliers and removed from the analysis (Neter, Kutner, Nachtsheim, & Wasserman, 1996). Regression coefficients were derived from bootstrapped estimates with 5000 replicates. Bayesian correlations (two-tailed, BF_10_) were conducted between VO_2_max and outcome variables for each multiple linear regression analysis. To assess the association between hippocampal structure and function, partial correlation analyses were also calculated between primary structural (left & right volume, NAA concentration) and functional (face-cued word recall and separation bias) measures, whilst accounting for the effect of age (Bonferroni-adjusted α-level = .008).

**Figure 3.**
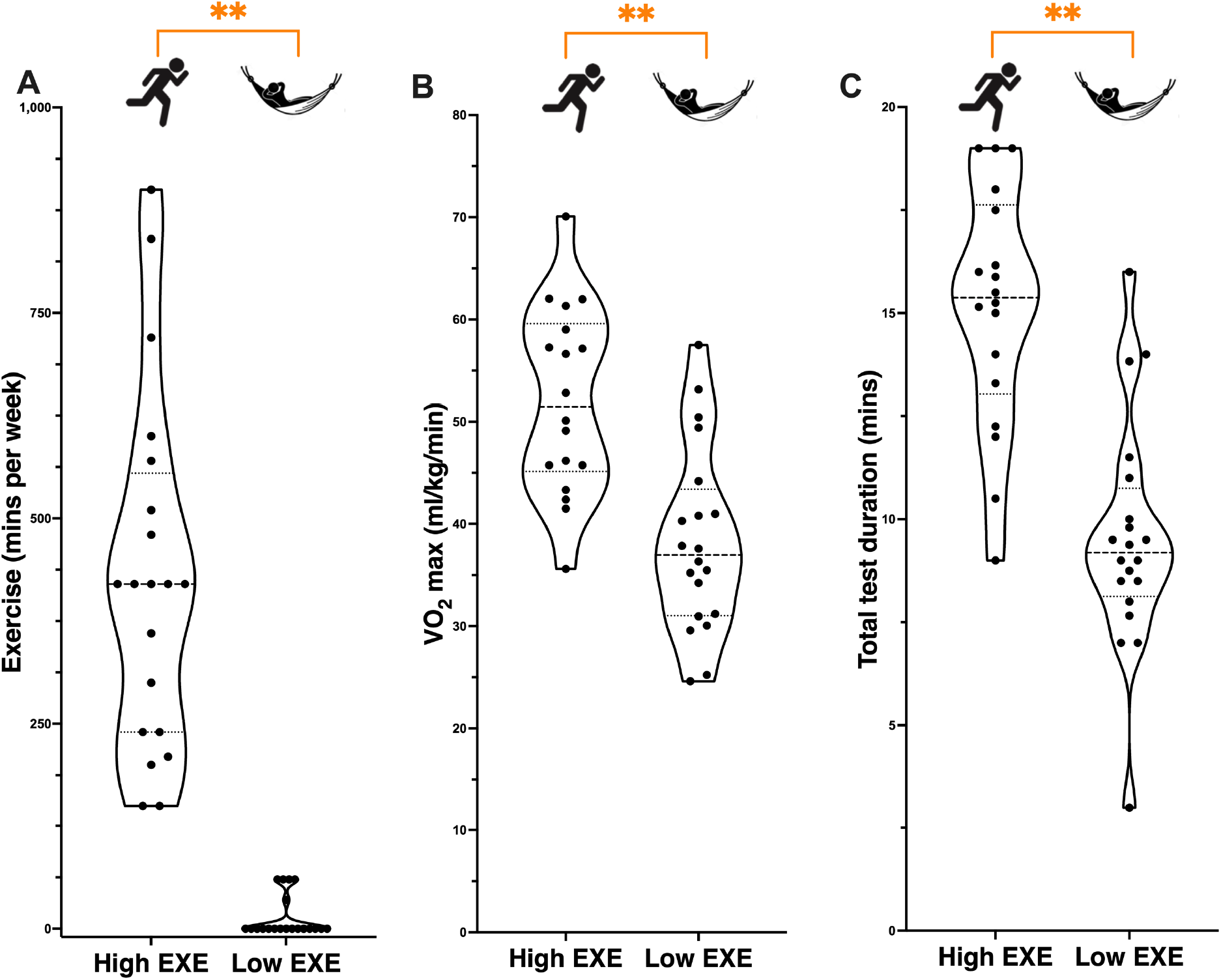
Comparison of self-reported weekly exercise and cardiorespiratory fitness between exercise (EXE) groups. (A) Self-reported weekly exercise. (B) Maximal oxygen consumption (VO_2_max) following a graded maximal exercise test. (C) Total test duration on graded exercise test. For all plots, filled circles indicate individual participants. For each group, mean values are indicated by horizontal dashed lines, whereas higher and lower quartiles are indicated by dotted lines. ** p < 5 × 10^−5^.

## Results

### Differences between EXE groups

There were no significant differences between EXE groups in regards to age (high EXE = 24.55 ± 8.77 years; low EXE = 26.40 ± 10.04 years, t(1,38) = −0.62, p = .54, Cohen’s d = 0.20), or sex (high EXE = 40 % female; low EXE = 65 % female, X^2^(1, N = 40) = 2.51, p = .11, Cramer’s V = .25).

We observed significantly higher VO_2_max levels, and total test durations (mins) on the graded exercise protocol, in the high EXE group relative to the low EXE group (see Table 1, Figure 3). Further, there was a significant positive association between self-reported weekly exercise and VO_2_max, with increasing weekly exercise minutes associated with greater CRF (r = .61, p = 4.29 × 10^−5^, supplementary figure S1).

**Table 1.**
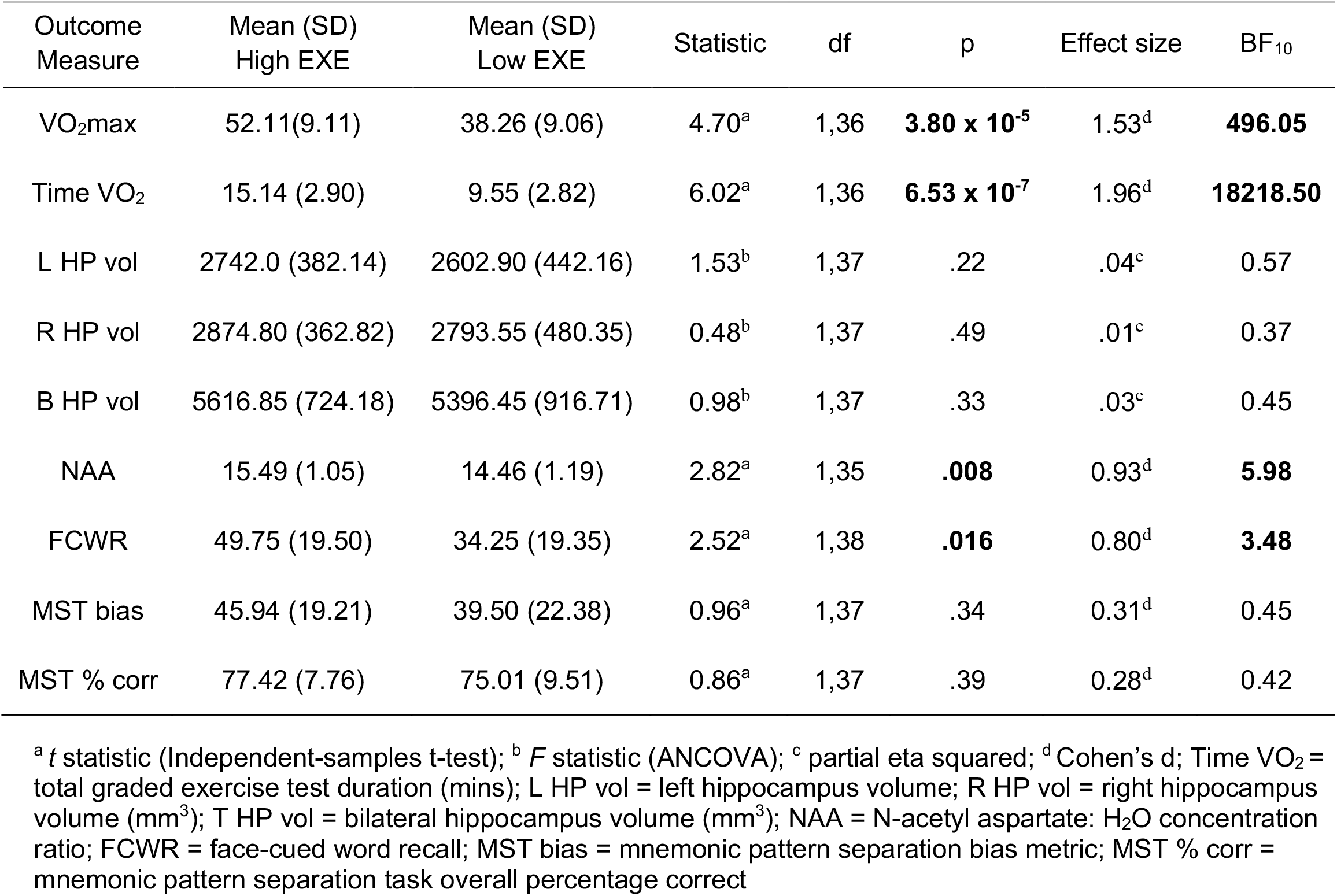
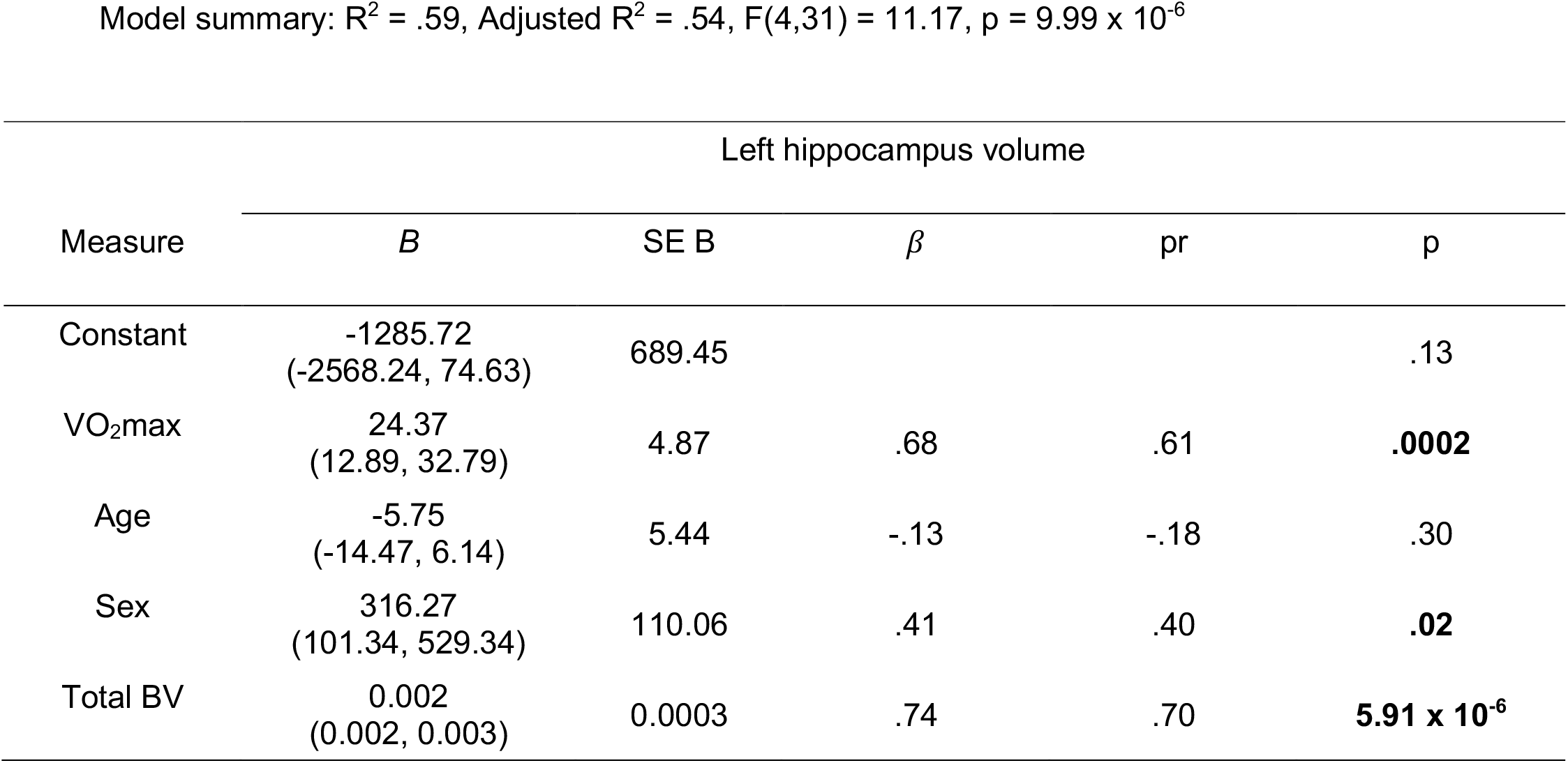
Analysis of fitness and hippocampal structure and function between High and Low EXE groups.

In regard to hippocampal structure and integrity, we did not find evidence for differences in left, right, or bilateral hippocampal volume between EXE groups (Table 1, Figure 4). Hippocampal volume estimates were positively skewed in the low EXE group (Figure 4). While ANCOVA models are generally considered robust to deviations from normality (Kéry & Hatfield, 2003; Levy, 1980; Olejnik & Algina, 1984), we conducted a supplementary analysis of left, right, and bilateral hippocampal volume with winsorization of the two most extreme values in the low EXE group. The results were unchanged for this supplementary analysis, with no significant differences in hippocampal volume between EXE groups (p > .05 for left, right, and bilateral hippocampal volume; see supplementary materials for complete analysis). However, we did observe significantly higher NAA:H_2_O concentration ratios in the High EXE group relative to the Low EXE group, with a BF_10_ of 5.98 indicating substantial evidence in favour of the alternative hypothesis (Table 1; Figure 5A; see supplementary materials for data quality analysis and comparison of grey matter and white matter voxel tissue fractions between EXE groups).

**Figure 5.**
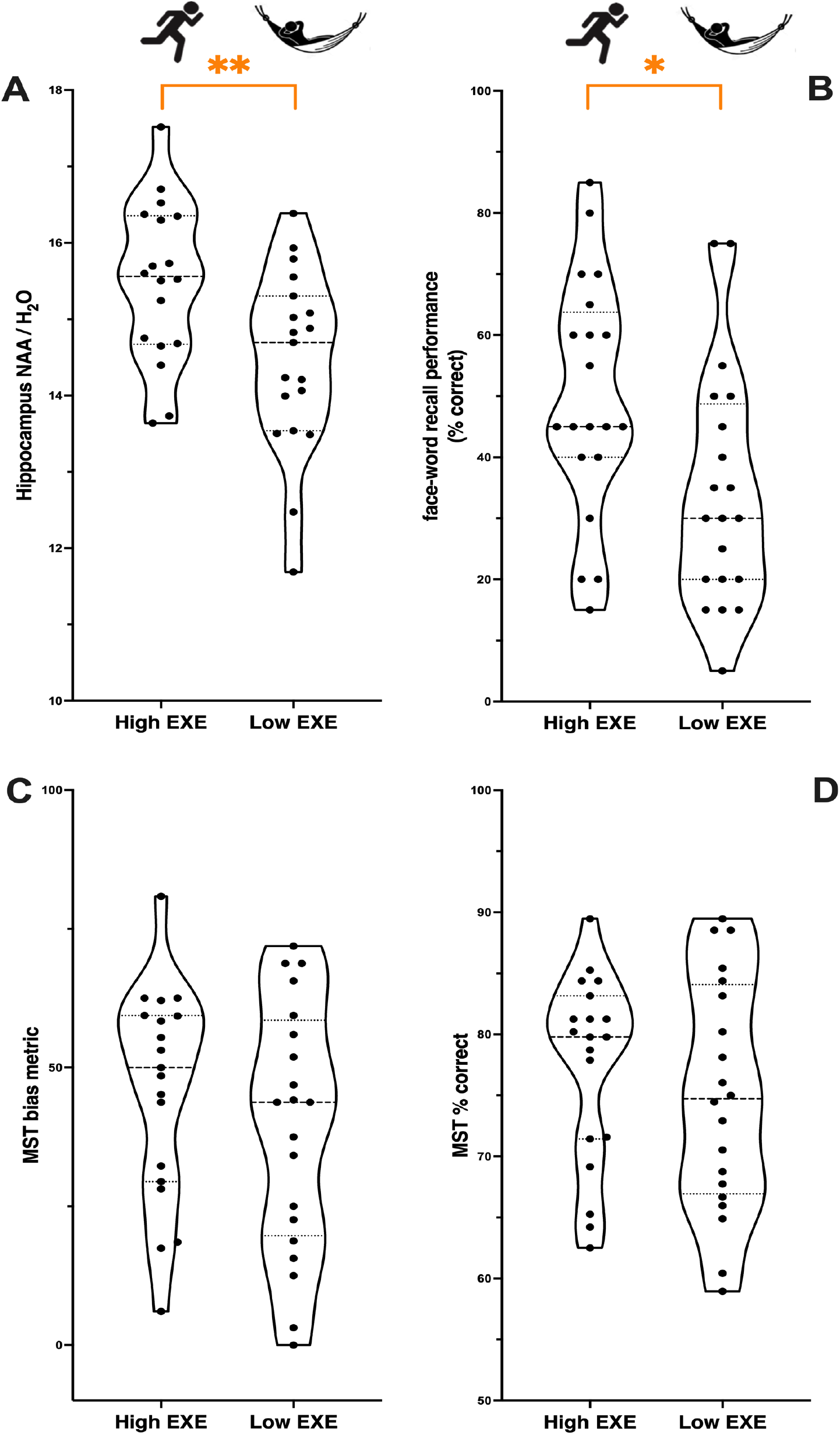
Comparison of NAA concentration and hippocampal function between EXE groups. **A)** Comparison of hippocampal N-acetyl-aspartate (NAA) concentration between EXE groups. **B)** Comparison of associative memory performance between EXE groups. **C-D)** Comparison of pattern separation (mnemonic similarity task; MST) performance between EXE groups. For all plots, filled circles indicate individual participants. For each group, mean values are indicated by horizontal dashed lines, whereas higher and lower quartiles are indicated by dotted lines. * p < .05; ** p < .01.

**Figure 4.**
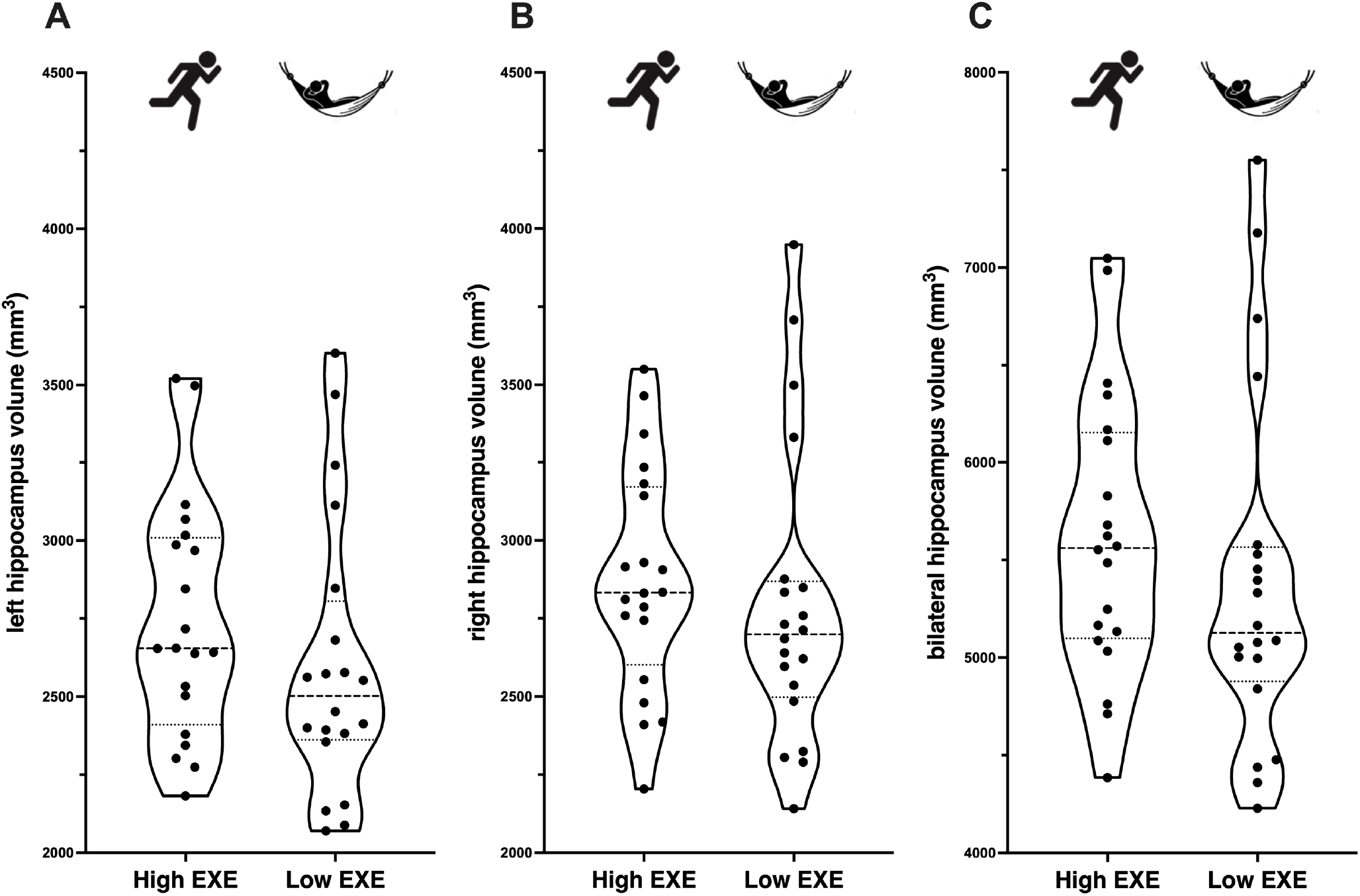
Comparison of hippocampal volume between EXE groups. Manually traced estimates of left **(A)**, right **(B)**, and bilateral **(C)** hippocampal volume were compared between EXE groups. There were no significant differences between EXE groups in left, right, or bilateral hippocampal volume. For all plots, filled circles indicate individual participants. For each group, mean values are indicated by horizontal dashed lines, whereas higher and lower quartiles are indicated by dotted lines.

In regard to hippocampal-dependent memory function, we observed significantly higher face-cued word recall performance in the high EXE group, relative to the low EXE group, with a BF_10_ of 3.48 indicating substantial evidence in favour of the alternative hypothesis (Table 1; Figure 5B). This finding remained significant when adjusting for multiple comparisons (Bonferroni-adjusted α-level = .017). However, we did not observe evidence of differences in pattern separation capacity between EXE groups (Table 1; Figure 5C-D). Together, these findings suggest that higher levels of EXE are associated with increased hippocampal NAA concentration and associative memory, but not hippocampal volume or pattern separation.

### Relationship between cardiorespiratory fitness and hippocampal structure and function

We assessed whether CRF level (VO_2_max) was positively associated with hippocampal volume. Two consistent outlier values (Cook’s distance > 3 × mean) were identified in each multiple linear regression model (i.e. left, right, bilateral volume analyses) and removed. There were significant positive associations between VO_2_max and hippocampal volume in the left hemisphere [t(1,35) = 4.32, p = .0002, BF_10_ = 2.38], right hemisphere [t(1,35) = 3.55, p = .001, BF_10_ = 1.47], and bilaterally [t(1,35) = 4.04, p = .0003, BF_10_ = 2.01] (Figure 6). VO_2_max uniquely explained 25% of the variance associated with left volume, 19% of right volume, and 23% of bilateral volume, and each of these associations remained significant when correcting for multiple comparisons (Bonferroni-adjusted α-level = .017) (see tables 1-3 in supplementary materials for model summary and regression coefficients). We next examined the relationship between CRF and hippocampal NAA concentration. One outlier value (Cook’s distance > 3 × mean) was identified and removed from the analysis. There was a significant positive association between VO_2_max and NAA concentration, t(1,34) = 3.39, p = .002, BF_10_ = 0.61 (Figure 7A), with VO_2_max uniquely explaining 23% of the variance in NAA concentration (see table 4 supplementary materials). At present, JASP does not allow categorical covariates to be included in Bayesian regression analyses (e.g. biological sex). Hence, the Bayes’ factor (BF_10_) was derived from the Bayesian bivariate correlation between VO_2_max and NAA concentration, without accounting for the variance associated with Age and Sex. Given that Sex was also a strong significant variable in the model (p = .0005), it is likely that the Bayes’ factor (BF_10_ = 0.61) has underestimated the strength of evidence in favour of the alternative hypothesis.

**Figure 6.**
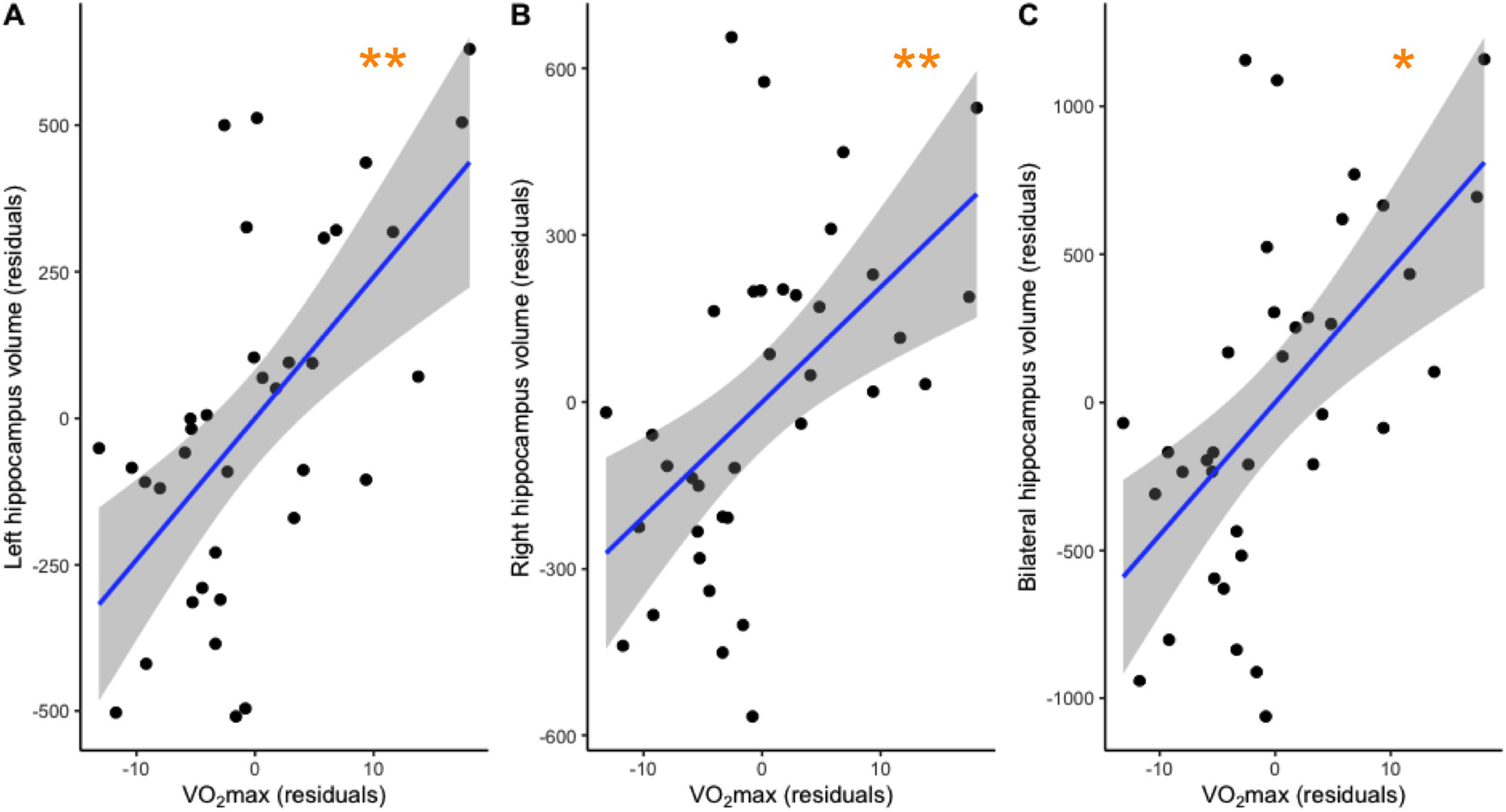
Associations between VO_2_max and hippocampal volume. There were significant positive associations observed between VO_2_max and left **(A)**, right **(B)**, and bilateral **(C)** hippocampal volume. Circles represent individual residual values, solid line shows regression line of best fit, grey shaded band depicts 95% confidence interval. Hippocampal volume residuals are represented on the y-axis and VO_2_max residuals are represented on the x-axis. * p <.01; ** p <.001.

**Figure 7.**
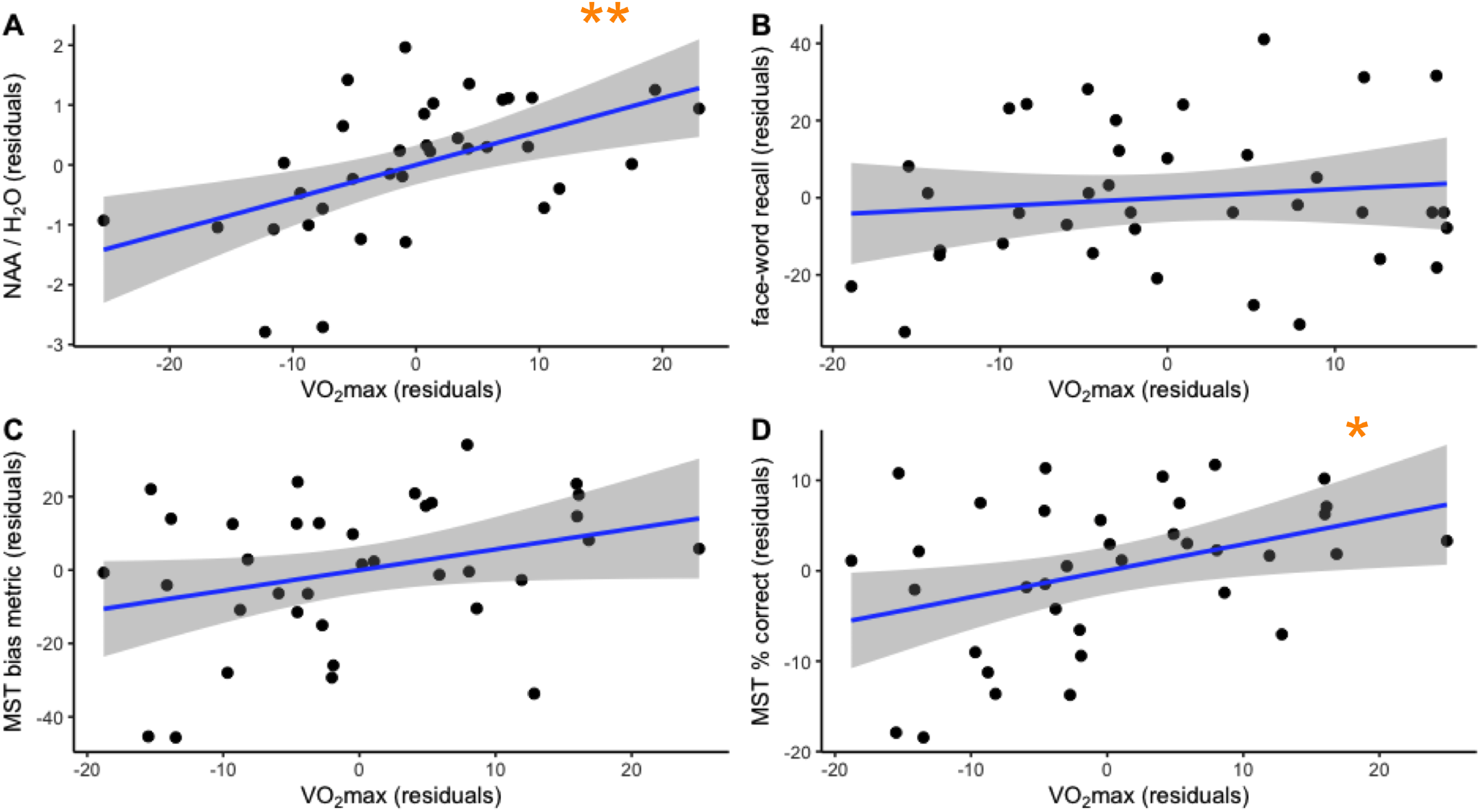
Associations between VO_2_max, NAA concentration, and hippocampal function. There were significant positive associations observed between VO_2_max and both NAA concentration and % correct on the mnemonic similarity task (MST). **(A)** Association between hippocampal NAA concentration and VO_2_max residuals. **(B)** Association between associative memory and VO_2_max residuals. **(C-D)** Association between pattern separation (**C** bias metric; **D** % correct) and VO_2_max residuals. Circles represent individual residual values, solid line shows regression line of best fit, grey shaded band depicts 95% confidence interval. Hippocampal measures are represented on the y-axis and VO_2_max is represented on the x-axis. * p <.05; ** p <.01.

Multiple linear regression models were also conducted to assess the relationship between CRF and face-cued word recall. One outlier value (Cook’s distance > 3 × mean) was identified in the face-cued word recall regression and removed. VO_2_max was not significantly associated with face-cued word recall performance (Figure 7B), t(1,34) = 0.71, p = .48, BF_10_ = 0.41, 1% variance explained (see table 5 in supplementary materials). The same regression model was also used to assess the relationship between VO_2_max and pattern separation performance. The outcome measures from the MST task (separation bias metric and overall percentage correct) were analysed separately. Bayesian analysis indicated there was anecdotal evidence in favour of the alternative hypothesis that VO_2_max was positively associated with separation bias (BF_10_ = 1.61) (Figure 7C), though this association did not exceed the significance threshold for the frequentist test, t(1,33) = 1.87, p = .070, 9% variance explained. There was also a significant positive association between VO_2_max and the overall percentage correct on the MST task, t(1,33) = 2.40, p = .022, (14% variance explained), with substantial evidence (BF_10_ = 4.09) in favour of the alternative hypothesis (Figure 7D). However, this association did not remain significant when correcting for multiple comparisons (Bonferroni-adjusted α-level = .017) (see tables 6-7 in supplementary materials). Further, while Bayesian analysis provided evidence in favour of the alternative hypothesis, including biological sex as an additional covariate in the frequentist regression model resulted in increased p-values (see tables 8–10 in supplementary materials).

### Relationship between hippocampal structure and function

We correlated hippocampal structural measures with each of the functional measures, controlling for the effect of age. There was a significant positive correlation between hippocampal NAA concentration and face-cued word recall performance (partial r = .46, p = .005, n = 36, see Figure 8). This association remained significant following correction for multiple comparisons (Bonferroni-adjusted α-level = .008). No other significant correlations were observed (all p values > .33).

**Figure 8.**
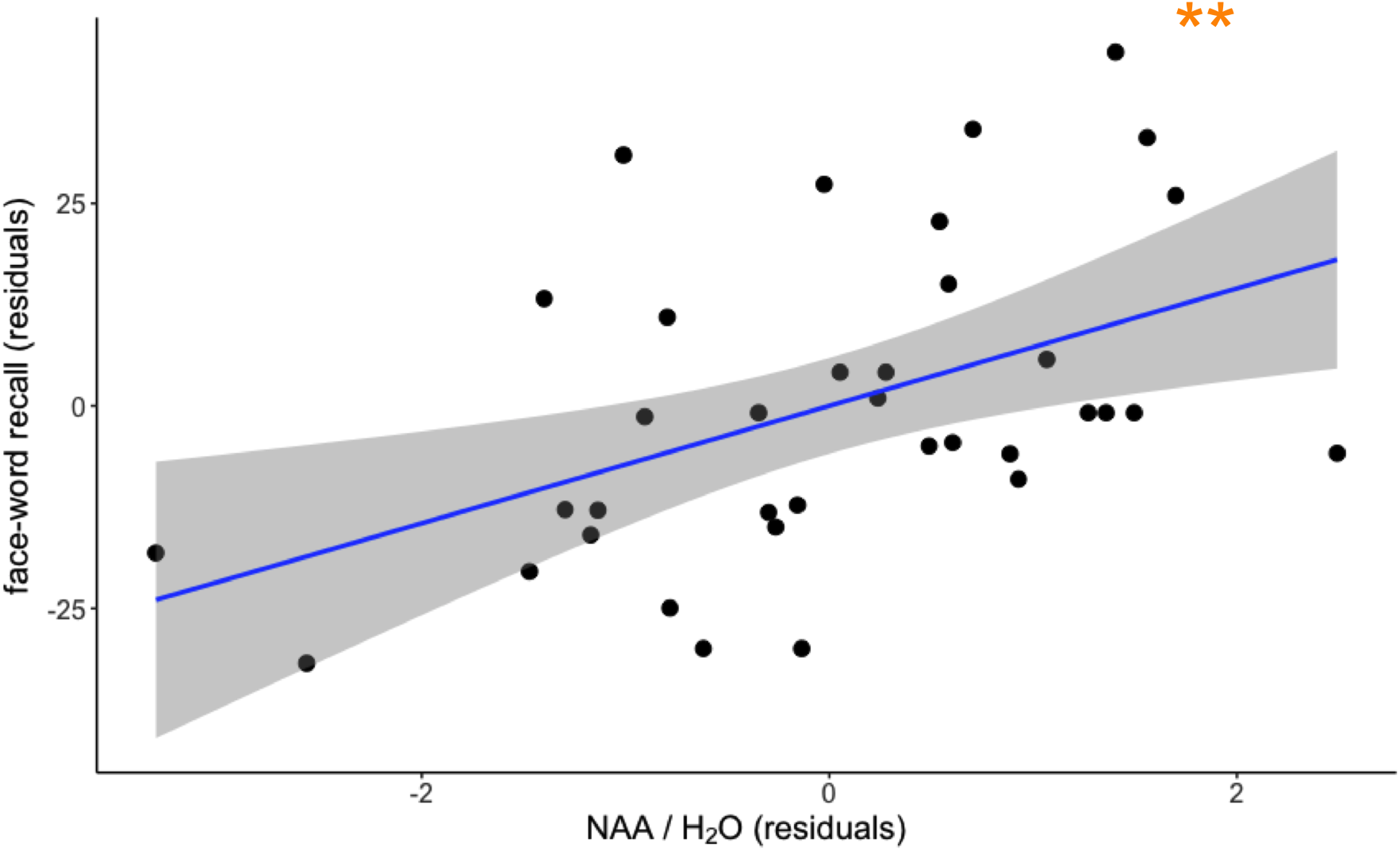
Associations between associative memory and NAA concentration. There was a significant positive association observed between associative memory and NAA concentration. Circles represent individual residual values, solid line shows regression line of best fit, grey shaded band depicts 95% confidence interval. Associative memory residuals are represented on the y-axis and NAA concentration residuals are represented on the x-axis. ** p <.01.

In summary, our findings suggest that higher CRF is associated with increased hippocampal volume, NAA concentration, and pattern separation capacity, but not face-cued word recall performance.

## Discussion

In this study, we investigated the associations of exercise and CRF with hippocampal structure and function in young to middle-aged adults. Using a multi-modal assessment, we compared hippocampal structure and function, and hippocampal-related memory performance between groups of individuals engaging in high versus low levels of exercise. We observed evidence of increased NAA concentration and associative memory performance in the high EXE group. However, no differences in hippocampal volume or pattern separation capacity were observed between EXE groups. We also investigated whether CRF (VO_2_max) was associated with measures of hippocampal structure and function. We found that CRF was positively associated with hippocampal volume (left, right, and bilateral), NAA concentration, and pattern separation, but not with associative memory. These findings demonstrate that regular exercise and cardiorespiratory fitness are consistently associated with greater hippocampal structure and function in young to middle age adults. These findings in this population are noteworthy because psychiatric illnesses that have been linked to hippocampal dysfunction are common in young and middle age adults (Kessler et al., 2005).

### Effects of exercise and cardiorespiratory fitness on hippocampal structure and integrity

Contrary to our hypothesis, we did not observe differences in hippocampal volume (left, right, or bilateral) between high and low EXE groups. Instead, using Bayesian statistics, we report anecdotal evidence in support of the null hypothesis. This finding is in contrast to a previous study by Killgore et al. (2013) which demonstrated a significant positive association between weekly exercise level and hippocampal grey matter volume in young to middle-aged adults. The discrepancy between study outcomes may be partially attributable to methodological/sampling differences. In the present study, we investigated differences in volume between individuals engaging in high vs low amounts of weekly exercise (i.e. meeting vs not meeting ACSM/WHO exercise guidelines). In contrast, Killgore et al., (2013) analysed associations between volume and exercise across a continuum of frequencies and durations (with 75% of individuals in their sample indicating some regular engagement in exercise). Therefore, it is possible that sampling a continuous distribution of exercise levels may be necessary to detect subtle changes to hippocampal volume related to recent exercise habits in these age groups. Indeed, a voxel-wise analysis of hippocampal regions of interest presented in the same study revealed no voxels surviving correction for multiple comparisons within the left hippocampus, and only a small cluster of 21 voxels surviving correction within an anterior portion of the right hippocampus (Killgore et al., 2013). Collectively, these findings suggest that the influence of current exercise engagement on hippocampal volume in young to middle-aged adults is likely subtle and requires further investigation in the context of well-powered experimental trial designs.

While we did not observe volumetric differences between EXE groups, we found that individual CRF level was positively associated with left, right, and bilateral hippocampal volume. This finding aligns with two studies that have reported similar relationships in young to middle-aged adults. Stillman et al. (2018) demonstrated an association between CRF and anterior hippocampal volume in a cross-sectional study of healthy young adults, and Pajonk et al. (2010) reported an association between CRF and total hippocampal volume increases following an exercise intervention in young to middle-aged adults with schizophrenia and an age-matched healthy control group (Pajonk et al., 2010). Thus, our reported relationship between CRF and hippocampal volume adds to a limited number of studies across early-to-mid-adulthood. The finding of a relationship between CRF and hippocampal volume is important as these age groups are over-represented in the diagnosis of psychiatric illnesses associated with hippocampal dysfunction. Therefore, promoting CRF may provide an effective means of ameliorating hippocampal function. Furthermore, a number of studies have shown that higher CRF prevents age-related hippocampal deterioration (Firth et al., 2017). Together, these findings indicate that higher CRF is associated with improved neuronal health across the adult lifespan, not just in older age.

The different pattern of results between CRF and self-reported exercise level may offer some insight into the timescales in which exercise interacts with hippocampal volume. Whilst there are multiple factors which contribute to individual differences in CRF (e.g. physiological/genetic predispositions), CRF also provides an indirect indicator of prior exercise history (Carrick-Ranson et al., 2014). An increase in CRF indicates physiological adaptation within the circulatory/respiratory systems in response to sustained, regular engagement in moderate/vigorous aerobic activity (which increase respiration and heart rate) (Caspersen, Powell, & Christenson, 1985). It is therefore possible that the positive relationship between CRF and hippocampal volume may reflect an accumulative effect of exercise. Namely, the repeated, sustained engagement in exercise required to demonstrate quantitatively high CRF may also be required for global changes to hippocampal volume. Engaging in aerobic exercise is known to upregulate a cascade of cellular and molecular factors within the hippocampus (e.g. BDNF, growth factors) which contribute to morphological effects (e.g. dendritic arborisation, synaptogenesis, vascularisation, and possibly neurogenesis) (Voss et al., 2013). It is possible that repeated triggering of these microscale mechanisms may be required before macroscale changes in hippocampal volume are detectable. Further, hippocampal blood flow is regulated by small, variably distributed arteries (ranging from 0.2 to 0.8 mm in diameter) branching off from the posterior cerebral and anterior choroidal arteries (Perosa et al., 2020; Spallazzi et al., 2019). It has been proposed that the effects of aerobic exercise on brain plasticity are primarily facilitated by angiogenesis and increased blood circulation (Stimpson, Davison, & Javadi, 2018). Given the variable and limited blood supply to the hippocampus, it is possible that increases in CRF (indicative of prolonged aerobic exercise over many weeks/months) are necessary for vascular plasticity and enduring structural changes within this region (Stimpson et al., 2018). Indeed, elevations in CRF following a 3-month aerobic exercise interventions are associated with increases in hippocampal blood flow, blood volume, and structural volume (Maass et al., 2015), suggesting that these mechanisms may respond over a longer timescale. While speculative, this explanation may also account for the absence of volumetric differences between EXE groups (categorised on the basis of current weekly exercise level rather than longer-term exercise history). Future studies are required to characterise this dose-response relationship between exercise and hippocampal volume in young to middle-aged adults.

In line with our expectations, we observed evidence of increased hippocampal NAA concentration in the context of both higher weekly exercise levels and CRF. Our results are consistent with a study by Pajonk et al. (2010) which reported elevated hippocampal NAA concentration following an exercise intervention in young to middle-aged adults with schizophrenia/age-matched healthy controls. Interestingly, higher CRF also offsets age-related NAA decline in the prefrontal cortex, and prefrontal NAA concentration mediates the positive association between CRF and working memory capacity (Erickson et al., 2012). NAA is predominantly localised to the cell body of neurons, where it plays a multi-factorial role in lipid synthesis/myelination and cell metabolism and is regarded as a marker of neuronal integrity/vitality (Moffett et al., 2007). Taken together, the results of our study and others indicate that higher CRF may confer neuroprotective effects in brain regions susceptible to age- and disease-related deterioration (i.e. hippocampus and prefrontal cortex). Furthermore, our results indicate enhanced NAA concentration in both high EXE individuals, and those with higher CRF. Given the cross-sectional nature of this study, future longitudinal studies are required to examine the timescale of these exercise-related effects.

### Effects of exercise and cardiorespiratory fitness on hippocampal function (associative memory, pattern separation)

Our findings provide some evidence of improved memory function in high EXE individuals. Bayesian analysis indicated substantial evidence of increased associative memory performance in individuals engaging in high levels of weekly exercise. However, no relationship between CRF and associative memory was found. Interestingly, past studies have found positive associations between CRF and associative memory level in young to middle-aged adults (Baym et al., 2014; Schwarb et al., 2017). Collectively these findings suggest that exercise is associated with improved associative memory performance in young to middle-aged adults. The observed association between NAA and associative memory in this study also suggests these effects may relate to hippocampal neuronal vitality/integrity. However, it is unclear what factors have contributed to the discrepancies between study outcomes in relation to CRF and associative memory. It is possible that differences in associative memory task demands may have contributed to observed inconsistencies, with our study assessing associative memory via face-word pairs, while past studies have utilised associative memory tasks with spatial memory and scene recognition components (Baym et al., 2014; Schwarb et al., 2017). Retrieval of associative memories involving faces vs spatial positions is known to activate a topographically distinct network of brain regions (Khader, Burke, Bien, Ranganath, & Rösler, 2005). Specifically, during retrieval of associative memories involving faces, several regions of the ventral visual pathway are preferentially activated (e.g. posterior fusiform gyrus/posterior cingulate). In comparison, associative spatial memory recall evokes stronger activation across dorsal visual regions (e.g. parietal cortex, dorsolateral prefrontal cortex, and precentral gyrus). In this sense, current exercise levels and CRF may preferentially affect certain forms of associative memory (e.g. those with spatial component), though future research is required to confirm this hypothesis.

In contrast, our results suggest that pattern separation capacity is positively associated with CRF, but not current weekly exercise level. However, we note that inclusion of biological sex as an additional covariate in these analyses weakened the observed associations. Nevertheless, this finding is in line with Déry et al. (2013) who reported a correlated improvement in CRF and pattern separation capacity following an exercise intervention, using a high-intensity interval training protocol (periodic running at 95% of age-predicted heart rate maximum, i.e. sprinting). In animals, exercise-induced improvements in pattern separation capacity are specifically linked to enhanced neurogenesis within the dentate gyrus (Clark, Brzezinska, Puchalski, Krone, & Rhodes, 2009; Creer, Romberg, Saksida, Van Praag, & Bussey, 2010). Increased BDNF expression is an important determinant of hippocampal neurogenesis (Rossi et al., 2006) and synaptic plasticity (Vaynman, Ying, & Gomez-Pinilla, 2004), and is elevated by exercise in an intensity-dependent manner (Saucedo Marquez, Vanaudenaerde, Troosters, & Wenderoth, 2015). While evidence supporting the existence of neurogenesis in adult humans remains equivocal, it is possible that increases in CRF (from regular moderate/high intensity exercise) are required to promote neurogenesis, thereby improving pattern separation in young/middle age groups.

### Future directions and conclusion

In this study, we highlight the positive associations between exercise and hippocampal structure and function in young to middle-aged adults. However, there are certain methodological limitations which should be acknowledged. Firstly, we utilised a cross-sectional design to compare hippocampal integrity between individuals engaging in high vs low levels of exercise. Whilst we conducted a multi-modal characterisation of hippocampal structure and function, we are unable to draw causal links between exercise and observed outcomes. Future studies are encouraged to evaluate the replicability of these findings in the context of longitudinal, controlled trials. Further, we have reported evidence linking both higher weekly exercise engagement and individual CRF to improved hippocampal health. The measurement of both self-reported weekly exercise level and individual CRF has allowed an assessment of their relative associations with improved hippocampal integrity. However, we did not collect a detailed history of each participant’s exercise engagement and thus the dose-response relationship between exercise and hippocampal outcomes cannot be determined. Lastly, in this study we focussed specifically on associations of exercise and CRF with the hippocampus. This decision was informed by a considerable evidence-base in older adults demonstrating that this region is particularly responsive to the effects of exercise (Kandola et al., 2016). However, such an approach may have failed to detect similar associations across other brain regions. Indeed, there is some evidence that CRF is positively associated with NAA concentration in the prefrontal cortex (Erickson et al., 2012). Further research is required to assess broader associations between exercise and brain structure and function in young to middle-aged adults.

In summary, we have provided evidence that both exercise and CRF are positively associated with hippocampal structure and function in young to middle-aged adults. We demonstrate that high levels of exercise are associated with elevated hippocampal integrity, while higher CRF is associated with increases in both volume and integrity. Further, we show that high levels of exercise are also associated with enhanced associative memory function. These findings have important implications for our understanding of how exercise and CRF relates to brain health in young to middle-aged adults. While a number of studies have demonstrated the capacity for exercise to mitigate deterioration in hippocampal function in older adults, we provide evidence that higher levels of exercise and CRF are associated with improved structure and function, prior to the expected onset of age-related deterioration (Raz, Rodrigue, Head, Kennedy, & Acker, 2004). Further, young and middle-aged adults are particularly vulnerable to psychiatric illnesses which affect hippocampal integrity. It is possible that exercise may provide a low-risk, effective method of remediating this dysfunction. Future work using longitudinal/interventional designs is required to develop a comprehensive understanding of the effects of exercise and cardiorespiratory fitness on brain health across the lifespan.

## Acknowledgements

We wish to thank the staff at Monash Biomedical Imaging for their assistance with MRI data acquisition, and Dr. Mark Mikkelson for technical support regarding NAA quantification within Gannet.

## Data and material availability

De-identified behavioural data are available at https://osf.io/gqrz8/. The conditions of our ethical approval do not permit open sharing of participant MRI data without prior informed consent. Therefore, we are unable to publicly archive the raw MRI data used in this study. Readers seeking access to the data should contact the lead author Joshua Hendrikse. Access will be granted to named individuals in accordance with ethical procedures governing the reuse of sensitive data. Specifically, requestors must complete a formal data sharing agreement and make clear the process by which participant consent would be sought. All code used for cognitive and MRI analysis is available at https://github.com/jhendrikse/ex_rtms_code.

## Supplementary materials

**Figure S1.**
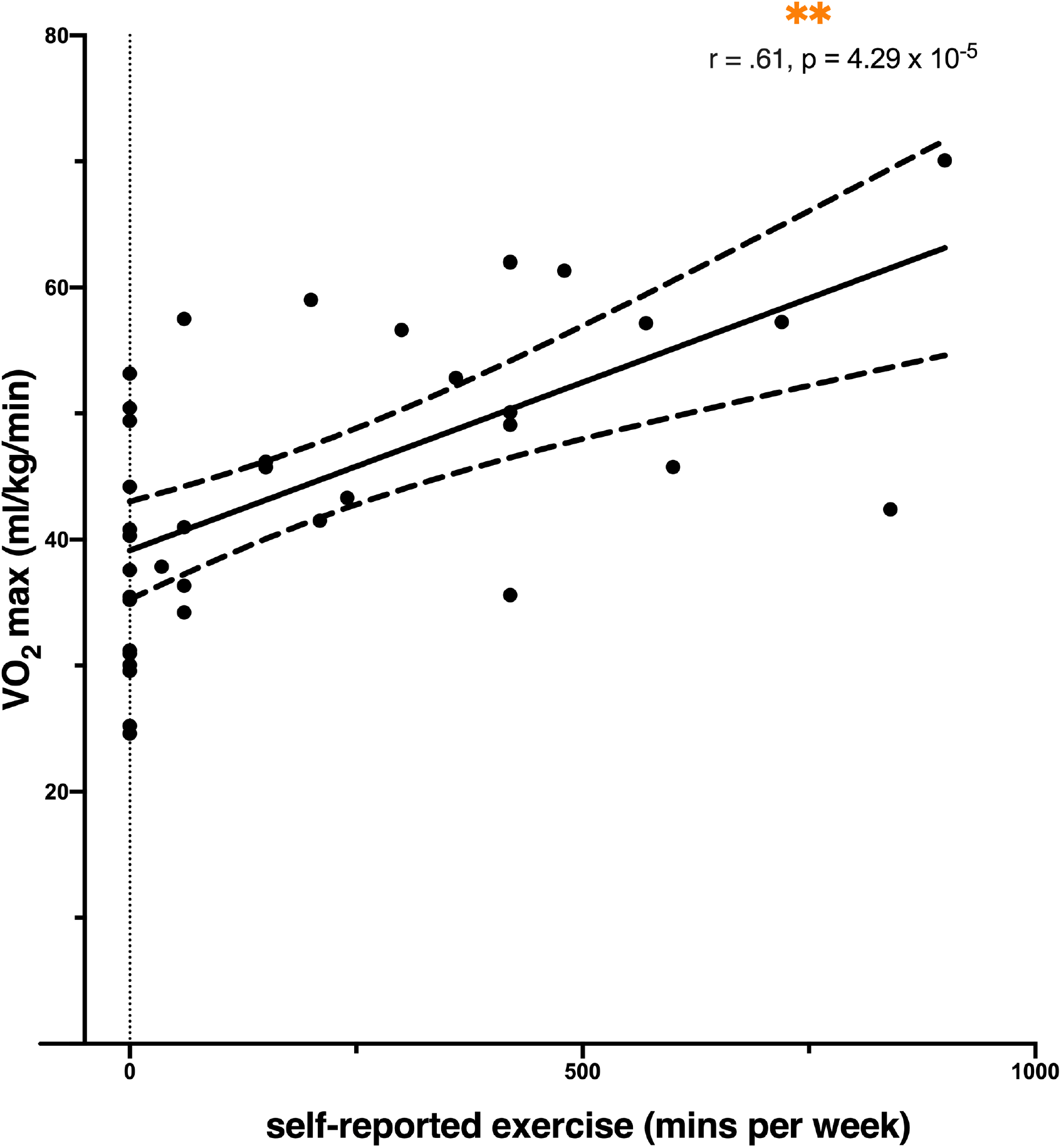
Association between self-reported exercise and VO_2_max. There was a significant positive association observed between self-reported exercise and VO_2_max (across EXE groups). Circles represent individual scores, solid line shows regression line of best fit, dotted lines depict 95% confidence interval. VO_2_max score is represented on the y-axis and exercise level (self-reported weekly minutes of exercise) is represented on the x-axis. ** p < 5 × 10^−5^.

**Figure S2.**
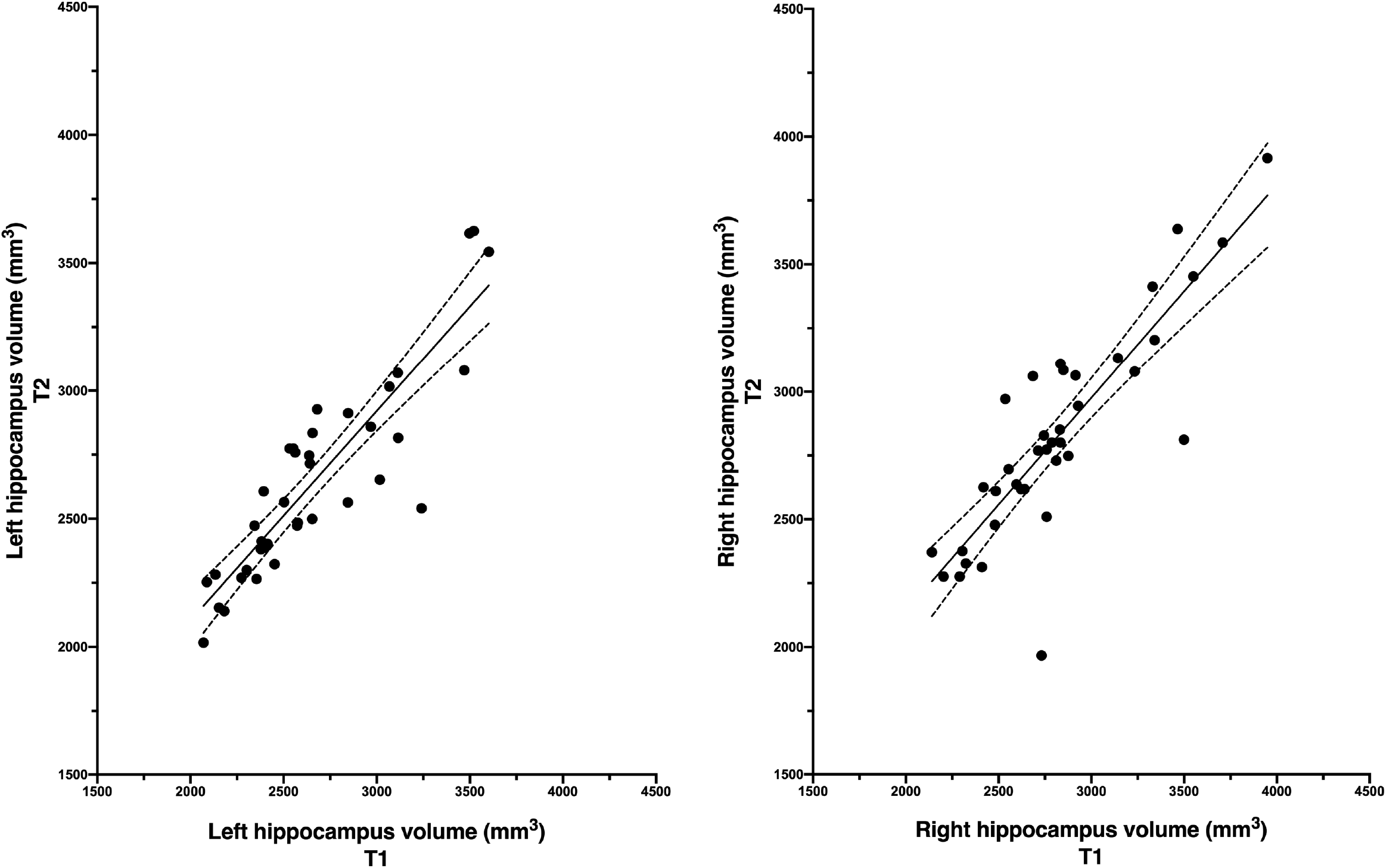
Intra-rater reliability of manual hippocampal tracing. To assess the intra-rater reliability of manual traced images, each subject’s hippocampi were retraced from a structural T1-weighted structural image acquired one week after the experimental assessment. T1-images acquired at this follow-up time point were acquired in the context of a separate study investigating the effects of four days of repetitive transcranial magnetic stimulation (rTMS) on functional connectivity (see Hendrikse et al., 2020). The tracer was blinded to MRI assessment time points and hypotheses pertaining to this study. To our knowledge, there is no evidence that similar rTMS protocols influence hippocampal volume in the healthy brain. Therefore, intra-rater correlation coefficients are unlikely to be biased by the intervening TMS application. The intra-rater correlation coefficient between experimental and follow-up assessments was .89 for left hippocampus and .86 for right hippocampus. The left plot depicts the association between manually traced estimates of left hippocampus volume at experimental time point (x-axis) and manually traced estimates of left hippocampus volume at the follow-up time point (y-axis). The right plot depicts the association between manually traced estimates of right hippocampus volume at experimental time point (x-axis) and manually traced estimates of right hippocampus volume at the follow-up time point (y-axis). Circles represent individual scores, solid line shows regression line of best fit, dotted lines depict 95% confidence interval.

### Comparison of manually traced estimates of hippocampal volume between experimental and follow-up assessments

To assess the intra-rater reliability of manual traced images, each subject’s hippocampi were retraced from a structural T1-weighted structural image acquired one week after the experimental assessment. To assess the consistency of hippocampal volume estimates across time points, separate 2 × 2 repeated-measures ANOVA was conducted on left and right hippocampal volume, with a within-subject factor of MRI Time Point (experimental, follow-up) and between-subject factor of EXE group (high EXE, low EXE). In regard to left hippocampal volume, there was no significant main effect of MRI Time Point (experimental, follow-up; *F*_1,35_ = 0.02, p = .90, *η*^2^_*p*_= .00), EXE Group (high EXE, low EXE; *F*_1,35_ = 2.47, p = .13, *η*^2^_*p*_ = .07), or significant Time Point × EXE Group interaction (*F*_1,35_ = 0.00, p = .97, *η*^2^_*p*_= .00). Similarly, in regard to right hippocampal volume, there was no significant main effect of MRI Time Point (experimental, follow-up; *F*_1,35_ = 1.48, p = .23, *η*^2^_*p*_= .04), EXE Group (high EXE, low EXE; *F*_1,35_ = 0.65, p = .43, *η*^2^_*p*_ = .02), or significant Time Point × EXE Group interaction (*F*_1,35_ = 0.03, p = .86, *η*^2^_*p*_= .00). Overall, there were no evidence of significant changes to total hippocampal volume in the week period between MRI assessments, suggesting that intra-rater reliability estimates are unlikely to be biased by systematic changes to hippocampal volume across MRI assessment time points.

### Analysis of sex-based differences of hippocampal-dependent memory

To account for possible sex-based differences in our measures of hippocampal function, we compared associative memory and pattern separation performance between males and females. There were no differences in associative memory performance [t(1,37) = −1.07, p = .29, Cohen’s d = 0.35], pattern separation bias [t(1,36) = 0.32, p = .75, Cohen’s d = 0.11], or pattern separation overall percentage correct [t(1,36) = 1.03, p = .31, Cohen’s d = 0.34] between groups, suggesting the observed outcomes are unlikely to have been influenced by biological sex.

### Analysis of NAA data quality

We assessed the consistency and reliability of NAA estimates between EXE groups. The assumption of normality was violated for NAA FWHM, water FWHM, and ENAA estimates. Therefore, these variables were compared between EXE groups using non-parametric Mann-Whitney tests. There were no significant differences in NAA FWHM between high EXE (M = 12.52, SD = 1.89) and low EXE (M = 13.14, SD = 4.38) groups, U = 192, p = .74, r = .07. Similarly, there were no differences in water FWHM between high EXE (M = 12.83, SD = 1.91) and low EXE (M = 12.63, SD = 1.44) groups, U = 182, p = .95, r = .01. There was also no difference in ENAA (high EXE M = 1.23, SD = 0.32; low EXE M = 1.29, SD = 0.39), U = 168, p = .74, r = .07, or water concentration between EXE groups (grand M = 1.181 × 10^−2^, grand SD = 9.3 × 10^−4^; high EXE M = 1.181 × 10^−2^, SD = 7.5 × 10^−4^; low EXE M = 1.181 × 10^−2^; SD = 1.08 × 10^−3^), t(1,36) = 0.02, p = .99, Cohen’s d = 0.01. Collectively, these results provide strong evidence of equivalent data quality and consistent NAA quantification between EXE groups. One outlier NAA:H2O value was identified (Z-score > 3.29, low EXE group) and removed from the analysis.

### Supplementary analysis of hippocampal volume between EXE groups

There was evidence of a positively skewed distribution of hippocampal volume estimates in the low EXE group (Skewness coefficient > 1, Shapiro-Wilk test p-value < .05). Thus, to explore the influence of this distribution on the reported outcomes, we conducted supplementary ANCOVAs on left, right, and bilateral hippocampal volume (dependent factor: Hippocampal Volume; fixed factor: EXE Group; covariate: Total Brain Volume) with removal of the two most extreme cases from the low EXE group. While there was a qualitative difference in hippocampal volume between groups following this manipulation, these effects were non-significant and do not alter the overall interpretation of results [left hippocampus, F(1,35) = 4.00, p = .053, s*η*^2^_*p*_ = 0.10; right hippocampus, F(1,35) = 2.49, p = .12, *η*^2^_*p*_ = 0.07; bilateral hippocampus, F(1,35) = 3,42, p = .073, *η*^2^_*p*_ = 0.09].

### Supplementary analysis of grey matter and white matter volume between EXE groups

We conducted partial volume segmentation of each subject’s T1-weighted anatomical image within each MRS voxel (figure 1) using FSL’s FAST. To explore possible differences in grey matter and white matter between EXE groups, ANCOVA’s were conducted separately on each tissue component (dependent variable: GM or WM volume; fixed factor: EXE Group; covariate: Total Brain Volume). There was evidence of greater grey matter volume in the low EXE group (M = 4078.59, SD = 547.22) relative to the high EXE group (M = 3660.01, SD = 461.35) [F(1,35) = 8.23, p = .007, *η*^2^_*p*_ = 0.19, BF_10_ = 6.57]. Conversely, we observed evidence of significantly greater white matter volume in the high EXE group (M = 4174.09, SD = 549.88) relative to the low EXE group (M = 3780.29, SD = 596.09) [F(1,35) = 4.84, p = .034, *η*^2^_*p*_ = 0.12, BF_10_ = 2.05]. Overall, these results provide evidence of differing tissue proportions between EXE groups within this region. However, given that the MRS voxel boundaries include non-hippocampal tissue, these estimates may not provide an accurate reflection of hippocampal tissue fractions. Further investigation with focal ROIs confined to the hippocampus is required to assess the validity of these findings.

**Tables 1-3:**
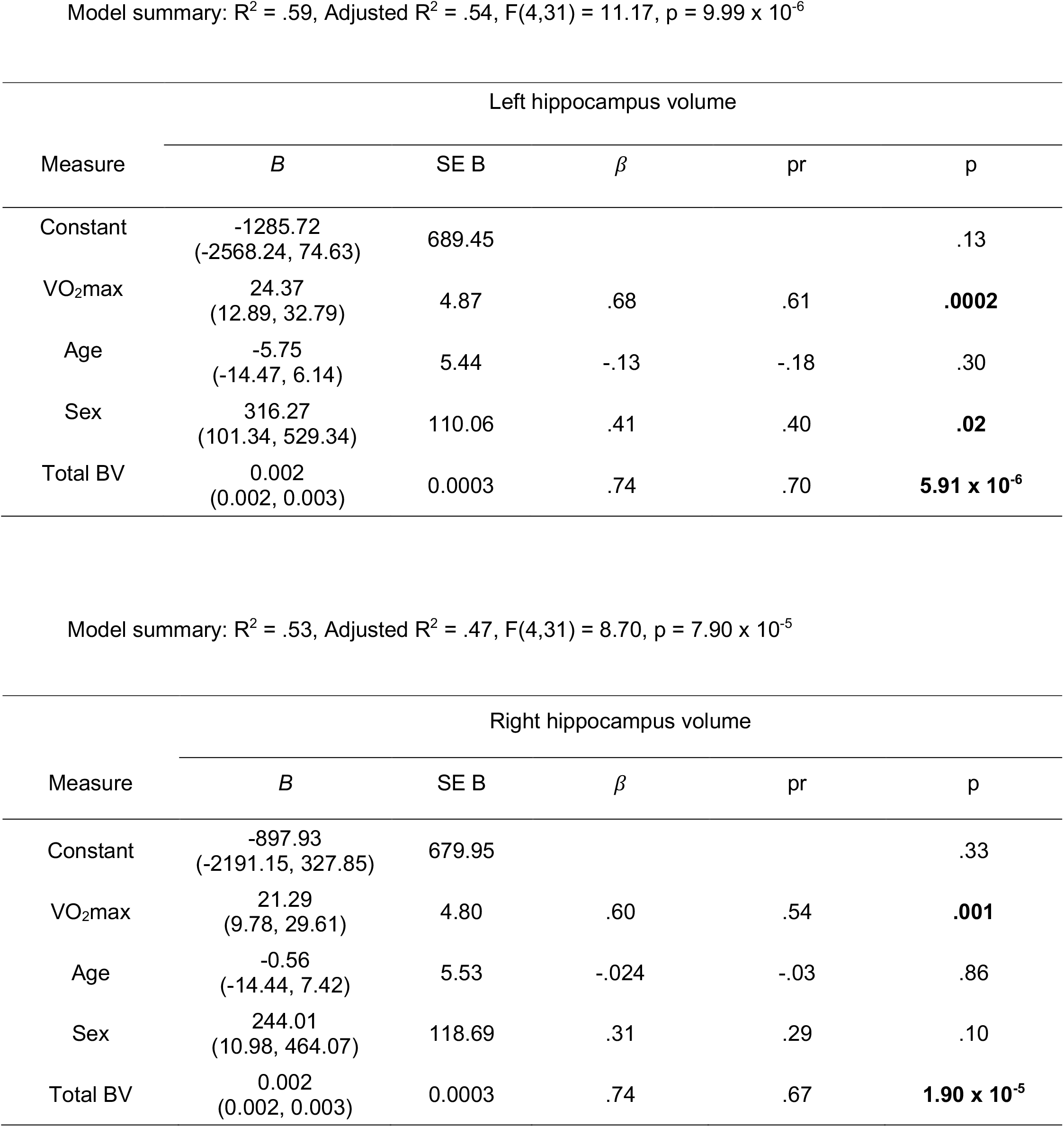
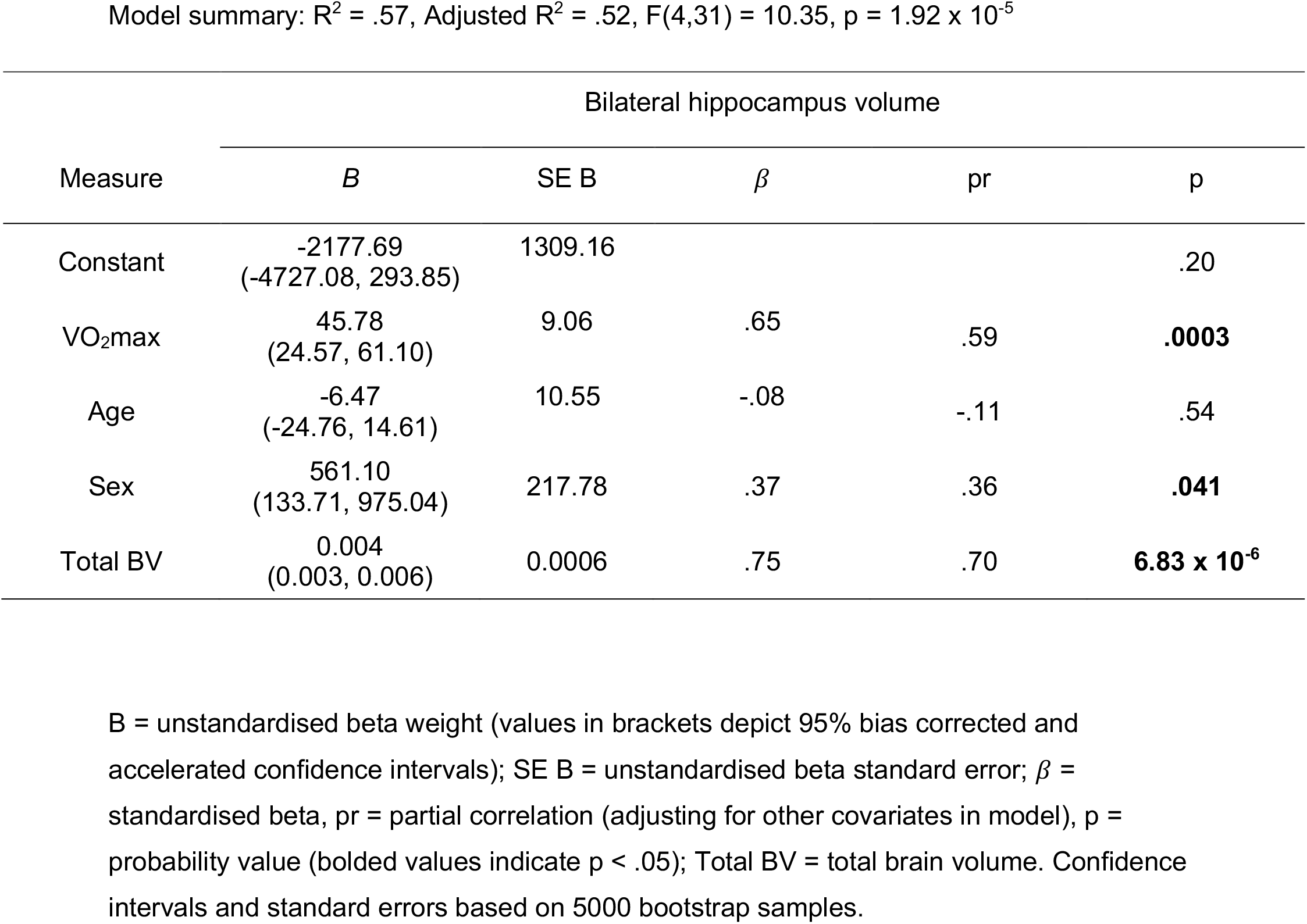
Regression statistics for hippocampus volume (DV) and VO_2_max, Age, Sex, Total Brain Volume (IVs)

**Table 4:** Regression statistics for hippocampus NAA concentration (DV) and VO_2_max, Age, Sex (IVs)

Model summary: $R^2 = .37$ , Adjusted $R^2 = .31$ , $F(3,31) = 6.12$ , $p = .002$ .
| Measure | Hippocampus NAA concentration |  |  |  |  |
| --- | --- | --- | --- | --- | --- |
| | *B* | SE B | $\beta$ | pr | p |
| Constant | 9.39<br>(5.14, 11.65) | 1.53 |  |  | **$3.36 \times 10^{-7}$** |
| VO<sub>2</sub>max | 0.056<br>(0.029, 0.096) | 0.016 | .55 | .52 | **.002** |
| Age | 0.029<br>(-0.010, 0.068) | 0.020 | .21 | .24 | .18 |
| Sex | 1.51<br>(0.86, 2.40) | 0.39 | .65 | .57 | **.0005** |

*B* = unstandardised beta weight (values in brackets depict 95% bias corrected and accelerated confidence intervals); SE B = unstandardised beta standard error; $\beta$ = standardised beta, pr = partial correlation (adjusting for other covariates in model), p = probability value (bolded values indicate $p < .05$ ). Confidence intervals and standard errors based on 5000 bootstrap samples.

### Examining the influence of grey and white matter tissue components on associations between CRF and hippocampal structure

We conducted supplementary analyses to explore the potential influence of inter-individual differences in grey and white matter tissue fractions on associations between CRF and hippocampal structure. Partial tissue fractions were derived from segmentation of the MRS voxel encompassing the left hippocampus. A multiple linear regression was conducted between left hippocampal volume and VO_2_max, with age, sex, total brain volume, grey matter, and white matter tissue fractions as additional covariates. This model did not explain additional variance when compared with the primary analysis [R^2^ = .59, Adjusted R^2^ = .50 with GM/WM covariates vs R^2^ = .59, Adjusted R^2^ = .54 without GM/WM], and a similar significant positive association was observed between VO_2_max and left hippocampal volume [t(1,35) = 4.07, p = .0004]. An additional multiple linear regression analysis was also conducted between NAA/water ratios and VO_2_max, with age, sex, grey matter, and white matter tissue fractions as additional covariates. The regression model with GM/WM covariates was comparable to the primary model [R^2^ = .38, Adjusted R^2^ = .28 with GM/WM covariates vs R^2^ = .37, Adjusted R^2^ = .31 without GM/WM], and a similar significant positive association was observed between VO_2_max and NAA concentration [t(1,34) = 3.01, p = .005]. In summary, the inclusion of these additional covariates did not significantly improve the model fit. These results indicate that variability in grey and white matter tissue fractions has not influenced the observed relationships between CRF and hippocampal structure and integrity.

**Table 5:** Regression statistics for face-cued word recall (DV) and VO_2_max, Age (IVs)

Model summary: $R^2 = .21$ , Adjusted $R^2 = .21$ , $F(2,34) = 4.54$ , $p = .018$ .
| Measure | Face-cued word recall |  |  |  |  |
| --- | --- | --- | --- | --- | --- |
| | *B* | SE B | $\beta$ | pr | p |
| Constant | 58.05<br>(20.60, 94.77) | 18.91 |  |  | **.003** |
| VO<sub>2</sub>max | 0.19<br>(-0.36, 0.88) | 0.31 | .11 | .12 | .48 |
| Age | -0.92<br>(-1.54, -0.26) | 0.32 | -.42 | -.42 | **.010** |

**Table 6-7:** Regression statistics for pattern separation (DV) and VO_2_max, Age (IVs)

Model summary: $R^2 = .16$ , Adjusted $R^2 = .11$ , $F(2,33) = 3.19$ , $p = .054$
| Measure | Separation bias metric |  |  |  |  |
| --- | --- | --- | --- | --- | --- |
| | *B* | SE B | $\beta$ | pr | p |
| Constant | 30.51<br>(-7.74, 68.87) | 19.53 |  |  | .085 |
| VO<sub>2</sub>max | 0.58<br>(-0.06, 1.19) | 0.32 | .30 | .31 | .070 |
| Age | -0.48<br>(-0.97, 0.37) | 0.33 | -.21 | -.22 | .20 |

Exercise and hippocampal structure & function Model summary: $R^2 = .19$ , Adjusted $R^2 = .14$ , $F(2,33) = 3.82$ , $p = .032$ .
| Measure | Overall percentage correct |  |  |  |  |
| --- | --- | --- | --- | --- | --- |
| | *B* | SE B | $\beta$ | pr | p |
| Constant | 66.86<br>(51.07, 83.79) | 8.42 |  |  | **$7.4 \times 10^{-11}$** |
| VO<sub>2</sub>max | 0.30<br>(0.06,0.56) | 0.13 | .38 | .39 | **.022** |
| Age | -0.13<br>(-0.32,0.17) | 0.12 | -.14 | -.15 | .39 |

B = unstandardised beta weight (values in brackets depict 95% bias corrected and accelerated confidence intervals); SE B = unstandardised beta standard error; $\beta$ = standardised beta, pr = partial correlation (adjusting for other covariates in model), p = probability value (bolded values indicate $p < .05$ ). Confidence intervals and standard errors based on 5000 bootstrap samples.

**Table 8:**
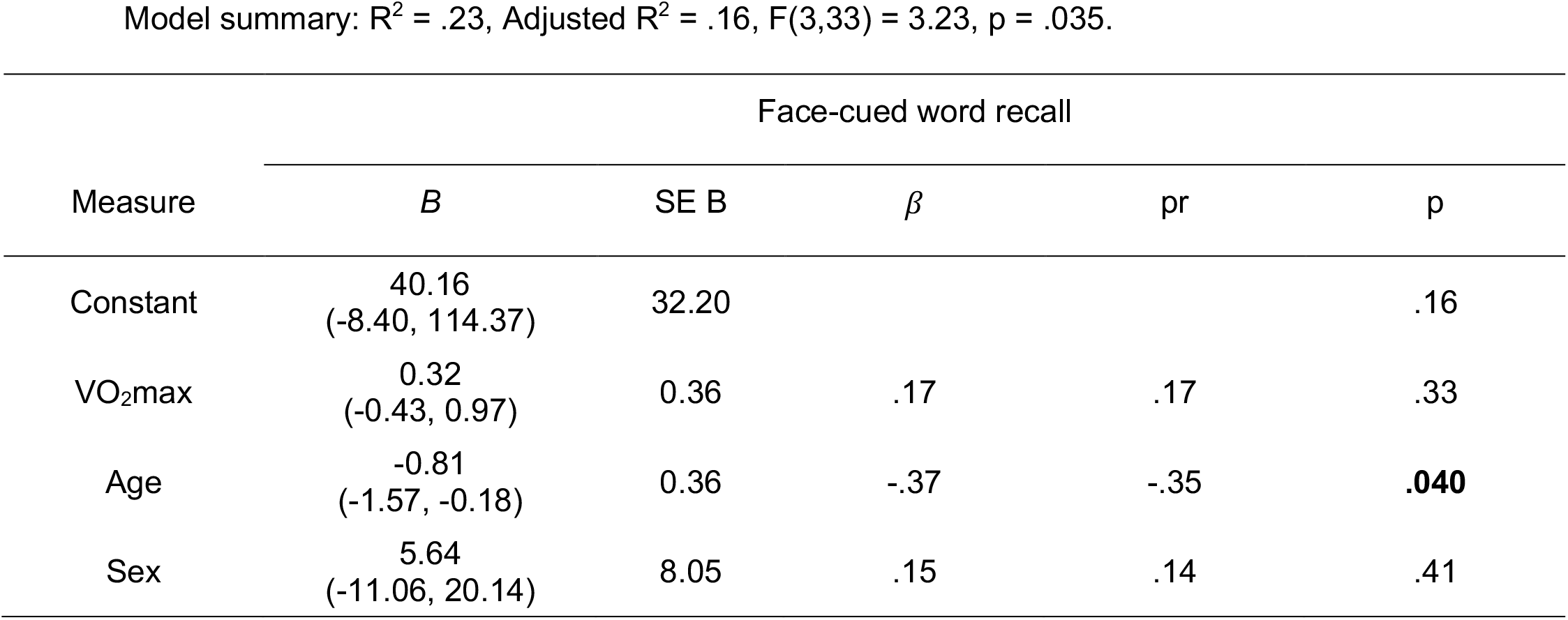
Regression statistics for supplementary analysis of face-cued word recall (DV) and VO_2_max, Age, Sex (IVs)

Model summary: $R^2 = .23$ , Adjusted $R^2 = .16$ , $F(3,33) = 3.23$ , $p = .035$ .
| Measure | Face-cued word recall |  |  |  |  |
| --- | --- | --- | --- | --- | --- |
| | *B* | SE B | $\beta$ | pr | p |
| Constant | 40.16<br>(-8.40, 114.37) | 32.20 |  |  | .16 |
| VO<sub>2</sub>max | 0.32<br>(-0.43, 0.97) | 0.36 | .17 | .17 | .33 |
| Age | -0.81<br>(-1.57, -0.18) | 0.36 | -.37 | -.35 | **.040** |
| Sex | 5.64<br>(-11.06, 20.14) | 8.05 | .15 | .14 | .41 |

**Table 9-10:** Regression statistics for supplementary analysis of pattern separation (DV) and VO_2_max, Age, Sex (IVs)

Model summary: $R^2 = .17$ , Adjusted $R^2 = .09$ , $F(3,32) = 2.14$ , $p = .11$ .
| Measure | Separation bias metric |  |  |  |  |
| --- | --- | --- | --- | --- | --- |
| | *B* | SE B | $\beta$ | pr | p |
| Constant | 41.13<br>(-7.28, 90.80) | 24.82 |  |  | .15 |
| VO<sub>2</sub>max | 0.50<br>(-0.15, 1.18) | 0.33 | .27 | .26 | .14 |
| Age | -0.55<br>(-1.11, 0.33) | 0.36 | -.24 | -.23 | .19 |
| Sex | -4.04<br>(-15.37, 10.09) | 6.40 | -.09 | -.08 | .65 |

Exercise and hippocampal structure & function Model summary: $R^2 = .20$ , Adjusted $R^2 = .13$ , $F(3,32) = 2.72$ , $p = .06$ .
| Measure | Overall percentage correct |  |  |  |  |
| --- | --- | --- | --- | --- | --- |
| | *B* | SE *B* | $\beta$ | pr | p |
| Constant | 74.34<br>(53.26, 94.12) | 10.43 |  |  | **$2.37 \times 10^{-7}$** |
| VO<sub>2</sub>max | 0.25<br>(-0.01, 0.52) | 0.13 | .33 | .31 | .07 |
| Age | -0.18<br>(-0.39, 0.15) | 0.14 | -.20 | -.19 | .27 |
| Sex | -2.59<br>(-7.26, 2.76) | 2.62 | -.14 | -.14 | .43 |

*B* = unstandardised beta weight (values in brackets depict 95% bias corrected and accelerated confidence intervals); SE *B* = unstandardised beta standard error; $\beta$ = standardised beta, pr = partial correlation (adjusting for other covariates in model), p = probability value (bolded values indicate $p < .05$ ). Confidence intervals and standard errors based on 5000 bootstrap samples.

